# Direct tissue sensing reprograms TLR4^+^ Tfh-like cells inflammatory profile in the joints of rheumatoid arthritis patients

**DOI:** 10.1101/2021.02.19.431679

**Authors:** Daniela Amaral-Silva, Rita C. Torrão, Rita Torres, Sandra Falcão, Maria João Gonçalves, Maria Paula Araújo, Maria José Martins, Carina Lopes, Agna Neto, José Marona, Tiago Costa, Walter Castelão, Ana Bento Silva, Inês Silva, Maria Helena Lourenço, Margarida Mateus, Nuno Pina Gonçalves, Santiago Manica, Manuela Costa, Fernando Pimentel-Santos, Ana Filipa Mourão, Jaime C. Branco, Helena Soares

## Abstract

CD4^+^ T cells mediate rheumatoid arthritis (RA) pathogenesis through both antibody-dependent and independent mechanisms. It remains unclear how synovial microenvironment impinges on CD4^+^ T cells pathogenic functions. Here, we identified a TLR4^+^ follicular helper T (Tfh) cell-like population present in the blood and expanded in synovial fluid. Mechanistically, we unveiled that homotypic T-T cell interactions through non-cognate HLA-DR:TCR contacts regulate TLR4 expression on T cells. TLR4^+^ T cells possess a two-pronged pathogenic activity. Upon TCR and ICOS engagement, TLR4^+^ T cells produce IL-21, a cytokine known to sponsor antibody production. However, direct TLR4^+^ engagement on T cells, by endogenous ligands in the arthritic joint, reprograms them towards an IL-17 inflammatory profile compatible with tissue damage program. Blocking TLR4 signaling with a specific inhibitor impaired IL-17 production in response to synovial fluid recognition. Ex vivo, synovial fluid TLR4^+^ T cells produced IL-17, but not IL-21. TLR4^+^ T cells appear to uniquely reconcile an ability to promote systemic antibody production with a local synovial driven tissue damage program. TLR4^+^ T cells could constitute an attractive cellular target and predictive biomarker for erosive arthritis.

## Introduction

In rheumatoid arthritis (RA) combined immune and joint tissue dysregulation synergize in propagating chronic inflammation and articular destruction. CD4^+^ T cells have been strongly implicated in RA pathogenesis through both antibody-dependent and independent mechanisms^1, 2^. It remains unclear, however, which CD4^+^ T cell population drives RA and how joint microenvironment impinges on their pathogenic functions. Unveiling CD4^+^ T cell pathogenic phenotype and its crosstalk with the arthritic joint environment would benefit diagnosis, patient stratification and could contribute to the design of better drugs that could effectively induce remission.

Effector functions sponsored by CD4 T cells in the joints constitute an active field of research. PD-1^high^CXCR5^high^ T follicular helper (Tfh) cells are the major T cell subset driving antibody production by B cells within secondary lymphoid organs^3, 4^. Even though circulating Tfh cell populations are diverse^5^, they have been defined as CXCR5^+ 6, 7^ and/or PD-1^+^CXCR5^+ 8^. In RA, various circulating Tfh cell populations have been correlated with B cell expansion and increased disease activity^9–11^. Recently, PD-1^+^CXCR5^-^ T cells, which share several markers with Tfh cells, were reported to infiltrate the inflamed synovium and to induce antibody production in vitro^12^. Notwithstanding, CD4^+^ T cell mediated antibody-independent mechanisms are at play in RA pathogenesis. Namely, IL-17 production by CD4^+^ T cells has been implicated in bone erosions^13, 14^ and cartilage damage^15–17^, with its neutralization reducing disease activity^18^ and curtailing cartilage and bone damage^13^. IL-17 production is regulated locally at the affected joint^19^, requiring both propitious tissue environment and cell-cell interactions, making it challenging to characterize IL-17 producing CD4^+^ T cells in RA. Thus, there is a pressing need to identify the synovium stimuli and the specific CD4^+^ T cell population responding to them driving IL-17 production.

T cell effector programs are profoundly shaped by the local tissue micro-environments where antigen recognition occurs^20^. RA joints are enriched in endogenous pro-inflammatory molecules and in pathogen recognition receptors that recognize them, namely Toll Like Receptors (TLRs). Polymorphisms in TLR4 have been found to be associated with increased RA susceptibility in humans^21^ and mice with TLR4 targeted deletions or loss-of-function mutations are protected from experimental arthritis^22–24^. In addition, TLR4 and its endogenous ligands are elevated in the synovial fluid and correlate with disease progression^23–27^. Even though predominantly expressed on innate immune cells, TLR4 has been found to be expressed at low levels in activated human and mice CD4 T cells^28, 29^. Curiously, TLR4 expression on T cells has been described to both facilitate and inhibit chronic inflammatory diseases^30^, with its pathological/protective role varying according to disease type and tissue affected. In a mouse model, TLR4 facilitates autoimmunity by functioning as a TCR co-receptor enhancing survival and proliferation, without affecting the type of quantity of cytokines produced^31^. It remains to be elucidated if TLR4 expression is enriched in CD4^+^ T cells of RA patients and whether the joint microenvironment engages TLRs directly on CD4^+^ T cells imprinting dysregulated inflammation and possibly diversifying their pathological function.

The strongest genetic association in RA is with HLA-DR alleles^32^. HLA-DR is constitutively expressed by antigen presenting cells (APCs) and interactions between antigen bearing HLA-DR on APCs and cognate TCR on CD4^+^ T cells drive full T cell activation^33^. Even though HLA-DR has been used as a marker of activated T cells for more than 40 years^34, 35^, whether or not HLA-DR expression plays a functional role on activated T cells has remained elusive.

We investigated the role of contextual cues in regulating T cell pathogenic programs in RA patients. We identified that RA patients possess a TLR4^+^ T cell population that is expanded in synovial fluid. Our data unveil that direct TLR4 stimulation on T cells goes beyond functioning as a coreceptor boosting TCR-driven response. Instead, TLR4 functions as a context sensor allowing to spatially tailor the pathological response elicited. Compatible with a systemic antibody response, TLR4^+^ T cells produce IL-21 upon TCR and ICOS engagement. Direct recognition of synovial components by TLR4 on T cells drives an inflammatory IL-17 program compatible with joint damage. Mechanistically, we uncovered for the first time a functional role for HLA-DR on T cells. We found that HLA-DR mediated homotypic T-T cell interactions regulate TLR4 expression on T cells, suggesting an important mechanism by which HLA-DR might drive RA disease susceptibility. Targeting the bidirectional communication between T cells and the joint tissue microenvironment might be critical to restore joint tissue homeostasis and induce RA remission.

## Results

### A circulating TLR4^+^CD4^+^ T cell population is expanded in the synovial fluid of RA patients

TLR4 is a robust tissue-damage sensor implicated in RA initiation and progression^25–27, 36, 37^. Previous studies focused on TLR4 expression by innate immune cells and synoviocytes^26, 27, 37^. Given CD4^+^ T cell role in RA and the abundance of TLR4 ligands in the arthritic joint, we investigated TLR4 expression by CD4^+^ T cells in freshly obtained synovial fluid from 9 RA patients (Table S1) undergoing arthrocentesis. Confirming our hypothesis, TLR4 was indeed expressed by ∼30% of CD4^+^ T cells in the synovial fluid (Fig. 1A). When compared to TLR4^-^ T cells, TLR4^+^ T cells, displayed bigger relative size and complexity, measured as FSC-A and SSC-A, respectively (Fig. 1B, C). Next, we assessed whether synovial fluid TLR4^+^ T cells would have a circulating counterpart by examining freshly obtained peripheral blood of 100 RA patients (Table S1). To ensure that we would be inclusive of CD4^+^ T cell with higher FSC-A/SSC-A, as the ones found in synovial TLR4^+^ T cells, we gated first on CD3^high^CD4^high^ T cells (Fig. 1B). Right away, we could detect two CD4^+^ T cell populations with distinct relative sizes and complexities. We performed doublet analysis, by plotting FSC-W versus FSC-A, we observed that these CD4^+^ T cell populations distribute along two distinct diagonals, suggesting that they are two distinct populations rather than cell conjugates. As determined for synovial TLR4^+^ T cells (Fig. 1A), TLR4 expression clustered on FSC-A^high^SSC-A^high^CD4^+^ T cells (Fig. 1D, E, F). The frequency of TLR4^+^ T cells in ranged between 0.02% and 28.7%, with a mean of ∼5% and mode of ∼1.3% (Fig. 1D). Donor matched analysis revealed a ∼4- and ∼12-fold enrichment in the frequency and expression levels, respectively, of TLR4^+^ T cells in synovial fluid relatively to the blood (Fig. 1 G, H). We found a correlation between the frequency of TLR4^+^ T cells in circulation and in the synovial fluid (Fig. 1I), suggesting that the blood faithfully reflects the enrichment of TLR4^+^ T cells in the synovial compartment.

**Figure 1.**
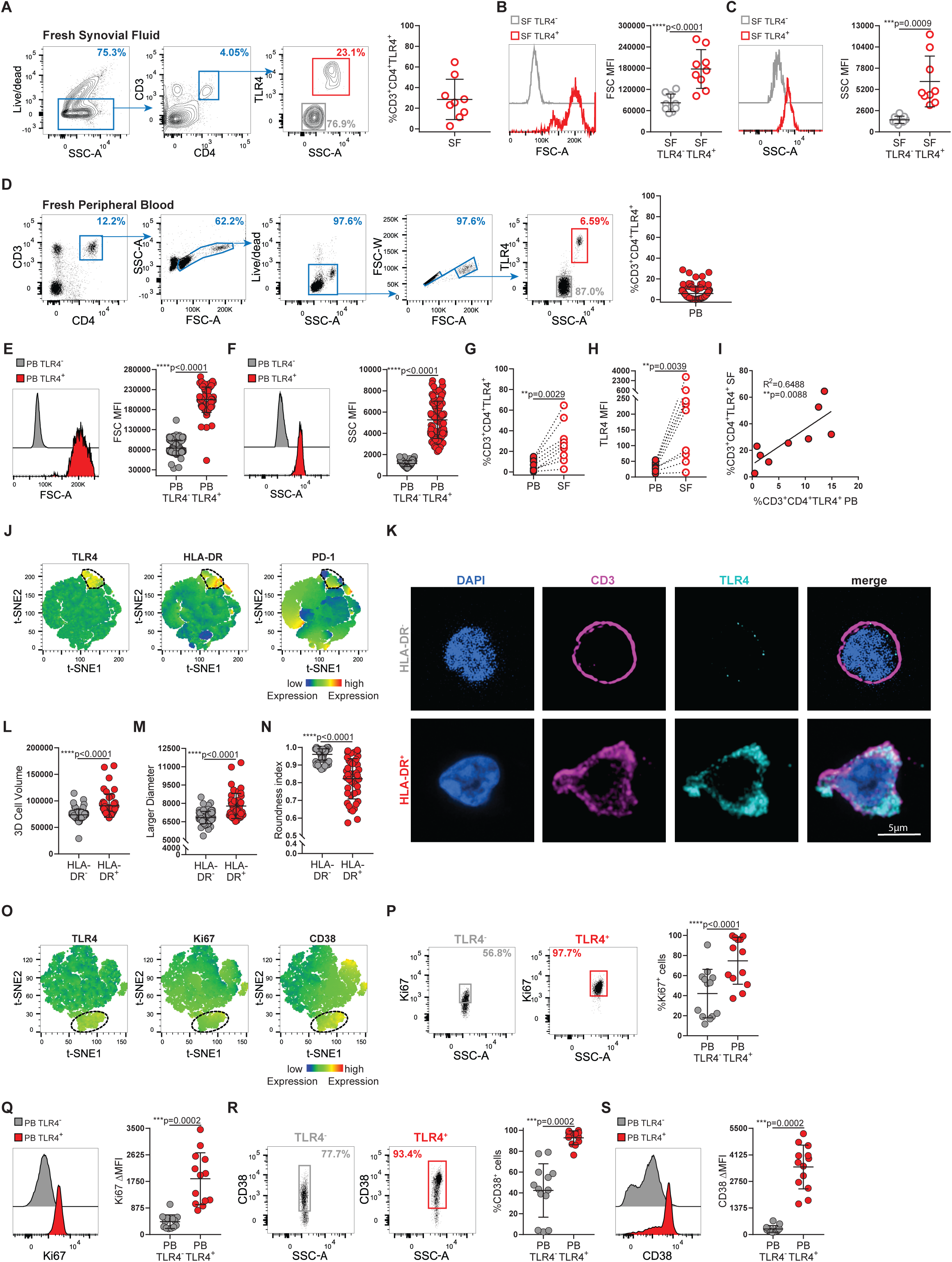
RA patients display a circulating TLR4^+^ T cell population that is expanded in the synovial fluid. (A) Gating strategy and cumulative frequency of CD3^+^CD4^+^TLR4^+^ cells in freshly obtained synovial fluid (n=9). (B) Representative histogram and cumulative plot of relative cell size (FSC-A) in TLR4^-^ (grey) and TLR4^+^ (red) synovial fluid T cells (n=9). (C) Representative histogram and cumulative plot of relative cell complexity (SSC-A) of TLR4^-^ (grey) and TLR4^+^ (red) synovial fluid T cells (n=9). (D) Gating strategy and cumulative frequency of CD3^+^CD4^+^TLR4^+^ cells in freshly obtained peripheral blood (n=100). (E) Representative histogram and cumulative plot of relative cell size (FSC-A) in TLR4^-^ (grey) and TLR4^+^ (red) peripheral blood T cells (n=100). (F) Representative histogram and cumulative plot of relative cell complexity (SSC-A) of TLR4^-^ (grey) and TLR4^+^ (red) peripheral blood T cells (n=100). (G) Donor matched analysis of the frequency of TLR4 expression by CD3^+^CD4^+^ T cells in peripheral blood (closed circles; PB) and in synovial fluid (open circles; SF) (n=9). (H) Donor matched analysis of the MFI of TLR4 expression by CD3^+^CD4^+^ T cells in peripheral blood (closed circles; PB) and in synovial fluid (open circles; SF) (n=9). (I) Correlation between the frequency of CD3^+^CD4^+^ TLR4^+^ T cells in blood (PB) and in synovial fluid (SF) (n=9). (J) t-SNE plots of peripheral blood total CD4^+^ T cells. Color indicates cell expression levels of labelled marker (TLR4, HLA-DR and PD-1). Circle demarks TLR4^+^ cells (n=26). (K-N) Confocal microscopy of FACS-purified HLA-DR^-^ and HLA-DR^+^ CD4^+^ T cells. (K) Cells were surface labelled for CD3 and TLR4, stained for DAPI and analyzed by 3D confocal microscopy. Bar, 5μm. (L) Cumulative graphs of 3D volume (M) larger diameter and (N) roundness index. (O) t-SNE plots of peripheral blood total CD3^+^CD4^+^ T cells. Color indicates cell expression levels of labelled marker (TLR4, Ki67 and CD38). Circle demarks TLR4^+^ cells. (P, Q) Representative dot plots and cumulative graphs of the frequency (P) and ΔMFI (Q) of Ki67 expression by TLR4^-^ and TLR4^+^ peripheral blood T cells (n=13 RA donors). (R, S) Representative dot plots and cumulative graphs of the frequency (R) and ΔMFI (S) of CD38 expression by TLR4^-^ and TLR4^+^ peripheral blood T cells (n=13). ΔMFI was calculated to correct for the distinct autofluorescence of the TLR4^-^ and TLR4^+^ T cell populations. ΔMFI was calculated by subtracting the fluorescence intensity minus one (FMO) from median fluorescence intensity (MFI) for each given marker. D’Agostino & Pearson normality test was performed. Shapiro-Wilk normality test was performed when n was too small for D’Agostino & Pearson normality testing. p values ****p<0.0001, ***p<0.001, **p<0.01, *p<0.05 were determined by (B, G, P) Paired t-test; (C, E, F, H, Q, R, S) Wilcoxon matched-pairs rank test; (I) Pearson Correlation and (L, M, N) Mann-Whitney test.

We reasoned that the increase in FSC-A and SSC-A values by synovial fluid and circulating TLR4^+^ T cells could reverberate their increased activation state. To address this possibility, we stained for T cell activation markers HLA-DR and PD-1. t-SNE analysis showed that PD-1 is expressed by various T cell populations, including TLR4^+^ T cells while, HLA-DR is selectively expressed by TLR4^+^ T cells (Fig. 1J). To formally exclude the possibility that bigger size of TLR4^+^ T cells was not due to cell aggregates and that a T cell population in RA patients does indeed express TLR4, we used HLA-DR as a proxy marker for TLR4^+^ T cells and sorted HLA-DR^+^ and HLA-DR^-^ CD4^+^ T cells by flow cytometry (Fig. S1A). We surface labelled HLA-DR^-^ and HLA-DR^+^ T cells for CD3 and TLR4 and analyzed them by confocal microscopy (Fig. 1K). Only, HLA-DR^+^ T cells displayed TLR4 at cell membrane, where it colocalized with CD3. The fact that TLR4 is evenly distributed throughout the cellular membrane disproves the possibility that these T cells gained TLR4 through trogocytosis^38^. As FSC-A only provides a relative measure of cell size, we calculated the 3D volume and measured larger width of both TLR4^-^ and TLR4^+^ T cells and found TLR4^+^ T cells to be bigger and wider than TLR4^-^ T cells (Fig. 1 K-M). Moreover, we observed that TLR4^+^ T cells exhibited membrane projections and alterations in their cell shape. To quantify the latter, we calculated roundness coefficient (ratio between the smallest and the larger diameter), where a roundness index of 1 characterizes perfectly round cells, with values <1 depicting a departure from it^33^. TLR4^+^ T cells roundness index was ∼0.8. Altogether, the higher FSC-A value of TLR4^+^ T cells is likely due to a combination of bigger cell size and alterations in cell shape caused by membrane projections.

TLR4 expression has been reported on senescent T cells in spondylarthritis patients^39^. Curiously, these TLR4^+^ senescent T cells were more prevalent in the blood than in the synovial fluid^39^. To exclude the possibility that the cells we identified are non-replicative senescent cells, we labelled them for the proliferation marker Ki67. We found TLR4^+^ T cells to be highly proliferative, with ∼75% of TLR4^+^ T cells undergoing cell cycle and ∼90% upregulating the activation marker CD38 (Fig. 1 O-S). Upregulation of HLA-DR, CD38 and Ki-67 by TLR4^+^ T cells supports their chronic activation, rather than a senescent, state. Since TLR4^-^ and TLR4^+^ have distinct cell sizes, they also display different autofluorescence. In order to correct for the effect of autofluorescence in our measurements we calculated ΔMFI by subtracting the fluorescence intensity minus one (FMO) from median fluorescence intensity (MFI) for each flow cytometer channel. We maintained this approach throughout all experiments. Collectively, we have identified a previously uncharacterized TLR4^+^ T cell population in RA patients. These TLR4^+^ T cells exhibit activation markers HLA-DR and CD38, a bigger cell size, and are highly proliferative, which is consistent with a T cell blast phenotype. Even though they can be detected in the blood, they are expanded in the synovial fluid, suggesting a role for these cells as drivers in RA pathology.

### TLR4^+^ T cell population correlates with anti-CCP antibody titers

Next, we pursued the relation between TLR4^+^ T cells and RA demographics, disease presentation and severity, and treatment. TLR4^+^ T cell frequency was not affected by age nor by biological gender (Fig. 2 A, B). RA has two clinical presentations, seropositive RA in which antibodies to either rheumatoid factor (RF) or to citrullinated (CCP) proteins are present and seronegative RA in which such antibodies are absent. TLR4^+^ T cells were present in both seropositive and seronegative patients (Fig. 2 C-E). Nonetheless, the frequency of TLR4^+^ T cells correlated with anti-CCP antibody titers in CCP^+^ patients (Fig. 2 F). The majority of patients in our cohort were either in clinical remission (66.7%) or presented low (12.3%) or moderate (18.5%) disease activity, with only 2.5% of the patients displaying a high disease activity score. Reflecting the high prevalence of patients with remitted or controlled disease (97.5%), we did not detect any correlation between disease activity score, measured either as DAS ESR (Fig. 2G) or DAS CRP (Fig. 2H) and TLR4^+^ T cells frequency (Fig. 2 G, H). Likewise, there was no detectable difference in TLR4^+^ T cell frequency when comparing different treatments by families (Fig. 2 I) nor for the supplementation of anti-inflammatory (Fig. 2 J) or corticosteroid (Fig. 2 K) drugs. When analyzed by individual drug use methotrexate (Fig. 2 L) and leflunomide (Fig. 2 M) exhibited a trend for slightly better and slightly worse outcomes, respectively, when compared to other drugs in the study (Fig. 2 L-P). Lastly, DMARD treatment duration does not seem to impact TLR4^+^ T cell frequency (Fig. 2 Q).

**Figure 2.**
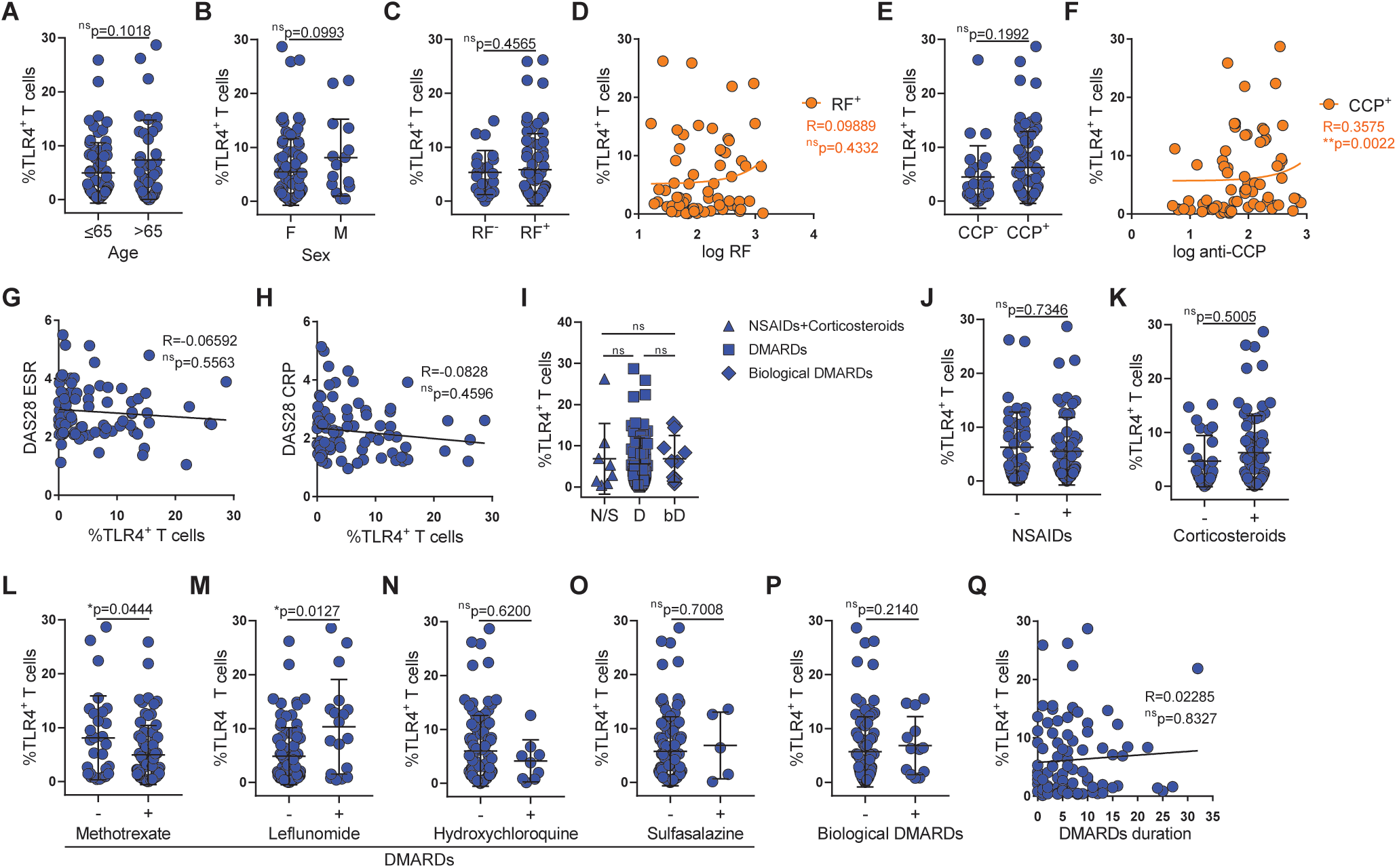
The frequency of TLR4^+^ T correlates with anti-CCP antibody titers and age, independently of treatment. (A) Frequency of TLR4^+^ T cells disaggregated by age (n=101; ≤65 years n=64; >65 years n=37). (B) Frequency of TLR4^+^ T cells disaggregated by sex (n=101; female n=86; male n=15). (C) Frequency of TLR4^+^ T cells disaggregated by factor rheumatoid (RF) status (n=88; RF^+^ n=66; RF^-^ n=22). (D) Correlation between factor rheumatoid titers and frequency of TLR4^+^ T cells in rheumatoid factor positive patients (n=66). (E) Frequency of TLR4^+^ T cells disaggregated by factor anti-CCP antibody status (n=96; CCP^+^ n=71; CCP^-^ n=25). (F) Correlation between factor anti-CCP antibody titers and frequency of TLR4^+^ T cells in CCP positive patients (n=71). (G) Correlation between frequency of TLR4^+^ T cells and DAS28 ESR score (n=83). (H) Correlation between frequency of TLR4^+^ T cells and DAS28 CRP score (n=83). (I) Frequency of TLR4^+^ T cells disaggregated by treatment family (N/S-NSAID and/or corticoids n=8; D-DMARDs n=81; bD-biological DMARDs n=12). (J-P) Frequency of TLR4^+^ T cells segregated by medication usage (n=101). (J) NSAIDs, (K) Corticosteroids, (L) Methotrexate, (M) Leflunomide, (N) Hydroxichloroquine, (O) Sulfasalazine, (P) biological DMARDs. (Q) Correlation between DMARD treatment duration and frequency of TLR4^+^ T cells (n=88). D’Agostino & Pearson normality test was performed. p values ****p<0.0001, ***p<0.001, **p<0.01, *p<0.05 were determined by (A, B, C, E, J, K, L, M, N, O, P) Mann-Whitney test; (D, F, G, H) Spearman Correlation and (I) Krustall-Wallis test with posttest Dunn’s multiple comparisons (N/S vs D and N/S vs bD ^ns^p>0.9999 and D vs bD ^ns^p=0.6963).

In summary, TLR4^+^ T cells persist in patients with controlled RA, regardless of treatment regimen, and correlate with anti-CCP antibody titers.

### HLA-DR mediated homotypic T-T cell interactions drive TLR4 surface expression

The strongest genetic association for developing RA is carried by HLA-DR alleles^32^. Even though HLA-DR has been used as a marker of T cell activation for more than 40 years, its functional role has remained elusive. Intrigued by the strong co-expression between HLA-DR and TLR4 (Fig. 1J, 3A), we analyzed the frequency of TLR4 expression by HLA-DR^+^CD4^+^ T cells (Fig. 3B) and reciprocally, the frequency of HLA-DR expression by TLR4^+^CD4^+^ T cells (Fig. 3C). While ∼80% of HLA-DR^+^CD4^+^ T cells co-expressed TLR4, ∼98% of TLR4^+^CD4^+^ T cells co-expressed HLA-DR. When looking at their cellular abundance, higher expression of HLA-DR was accompanied by greater TLR4 expression (Fig. 3D). Taken together, the above data suggested that there might be a link between HLA-DR and TLR4 expression.

**Figure 3.**
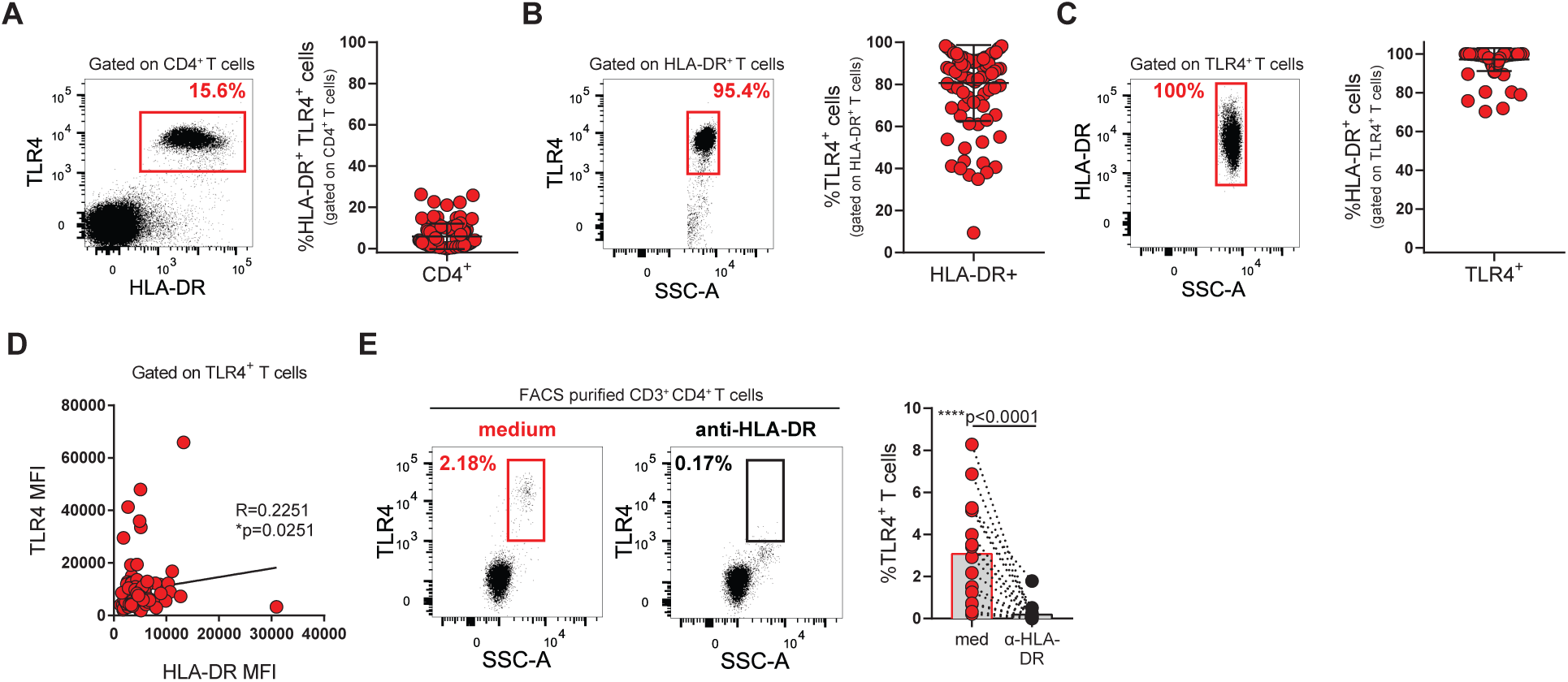
Blocking HLA-DR abrogates TLR4 surface expression in T cells. (A) Representative plots and cumulative graph (n=99) of the frequency of HLA-DR^+^TLR4^+^ T cells. (B) Representative plots and cumulative graph (n=99) of the frequency of TLR4 expression by HLA-DR^+^ T cells. (C) Representative plots and cumulative graph (n=99) of the frequency of HLA-DR expression by TLR4^+^ T cells. (D) Correlation between HLA-DR and TLR4 MFIs in TLR4^+^ T cells (n=99). (E) Representative plots and cumulative graph (n=17) of the frequency of TLR4^+^ T cells after incubating FACS-purified CD4^+^ T cells with a blocking antibody to HLA-DR for 18 hours. D’Agostino & Pearson normality test was performed. p values ****p<0.0001, ***p<0.001, **p<0.01, *p<0.05 were determined by (D) Spearman Correlation and (E) Wilcoxon matched-pairs rank test.

Recognition of noncognate-antigen:HLA-DR complexes on APCs by the TCR, albeit incapable of driving full T cell activation, generates nuanced effects on T cell activation and gene expression^40, 41^. We posited that homotypic T-T cell interactions through non-cognate HLA-DR:TCR contacts could control TLR4 expression on T cells. To address this possibility, we FACS-purified circulating CD4^+^ T cells with purity >99% (Fig. S1 B, C) and incubated them overnight with anti-HLA-DR blocking antibody or medium (Fig. 3E). Blocking HLA-DR dependent T-T cell contacts, led to a stark decrease in TLR4 surface expression (Fig. 3E). Indicating that HLA-DR regulates TLR4 expression through homotypic T-T cell interactions, in which HLA-DR on a T cell engages TCR on a neighboring one.

Altogether, our data identifies, for the first time, a functional role for HLA-DR on CD4^+^ T cells in which homotypic T-T cell interactions through HLA-DR:TCR contacts regulate TLR4 expression and suggest a novel mechanism by which HLA-DR might drive RA disease susceptibility.

### TLR4^+^ T cells share features of Tfh cells

Tfh-like T cells have been implicated in RA and other chronic inflammatory diseases due to their capability to induce antibody production^11, 12, 42^. We checked whether TLR4^+^ T cells would share Tfh features, namely high expression of chemokine receptor CXCR5 and of the co-receptors PD-1 (Fig. 1J) and ICOS. Even though CXCR5 (Fig. 4 A-C) and PD-1 (Fig. 4 A, D, E) could be detected in both TLR4^-^ and TLR4^+^ T cell populations, they were enriched in TLR4^+^ T cells with a co-expression of ∼80% (Fig. 4F). Curiously, ICOS was more expressed in TLR4^-^ than in TLR4^+^ T cells (Fig. 4 A, G, H). Nonetheless, in TLR4^+^ T cells co-expression of ICOS and CXCR5 (Fig. 4H) and ICOS and PD-1 (Fig. 4J) was enriched. The fact that TLR4^+^ T cells are enriched of CXCR5 and PD-1 suggests that they might consist a circulating Tfh-like population^6, 7^. To characterize this further, we explored whether the enrichment in TLR4^+^ T cells could reflect the frequency of circulating Tfh cells. TLR4^+^ T cell frequency positively correlated with the frequency of CXCR5^+^ (Fig. 2K) and PD-1^+^ (Fig. 2L) circulating CD4^+^ T cells.

**Figure 4.**
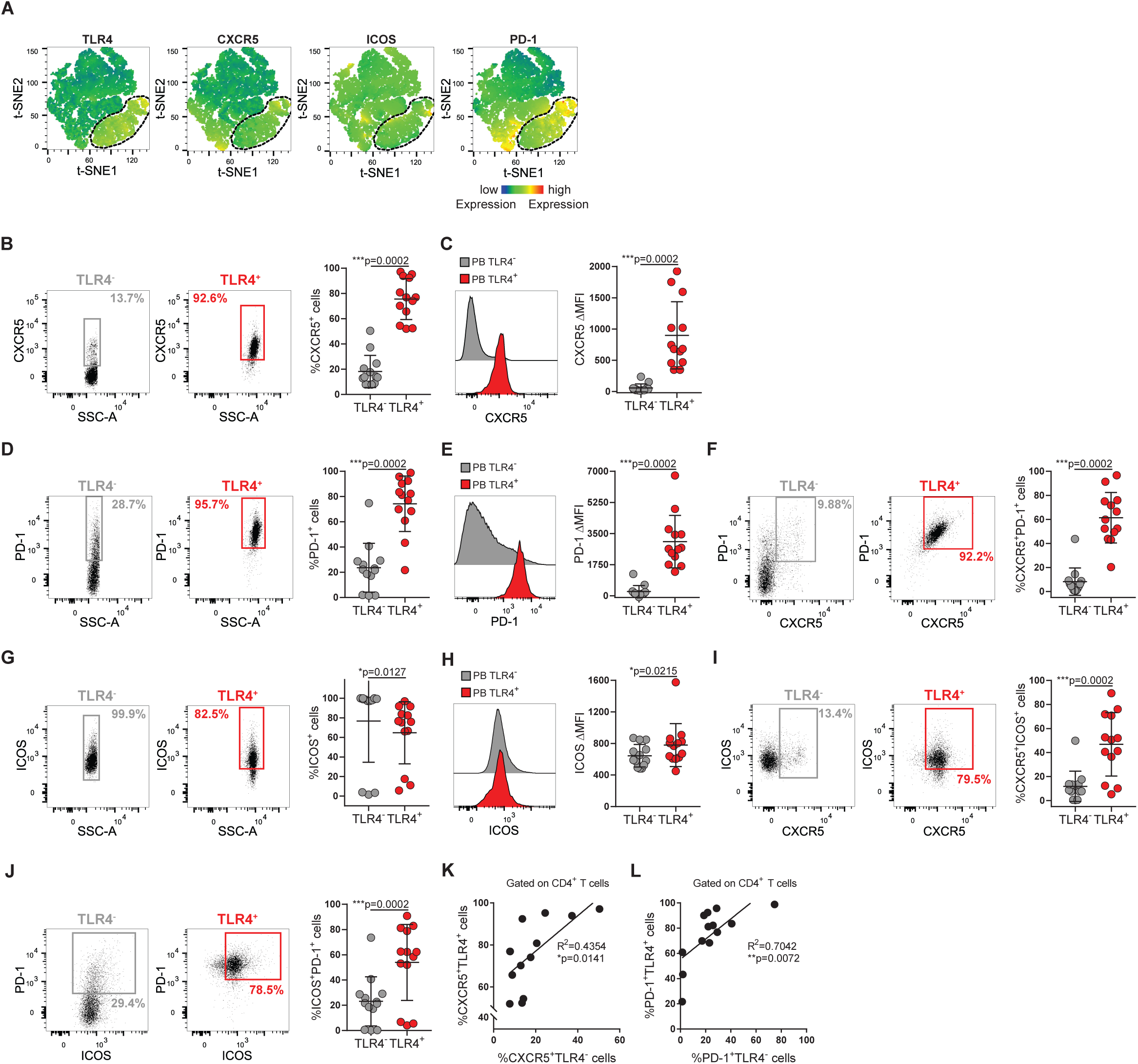
TLR4^+^ T cells have features of Tfh-like cells. (A) t-SNE plots of peripheral blood total CD4^+^ T cells. Color indicates cell expression levels of labelled marker (TLR4, CXCR5, ICOS and PD-1). Circle demarks TLR4^+^ cells. (B, C) Representative plots and cumulative analysis (n=13) of CXCR5 frequency (A) and *Δ*MFI (B) in TLR4^+^ (red) versus TLR4^-^ (grey) T cells. (D, E) Representative plots and cumulative analysis (n=13) of PD-1 frequency (D) and *Δ*MFI (E) in TLR4^+^ (red) versus TLR4^-^ T cells (grey). (F) Representative plots and cumulative analysis (n=13) of the frequency of CXCR5 and PD-1 co-expression TLR4^+^ (red) versus TLR4^-^ (grey) T cells. (G, H) Representative plots and cumulative analysis (n=13) of ICOS frequency (G) and *Δ*MFI (H) in TLR4^+^ (red) versus TLR4^-^ (grey) T cells. (I) Representative plots and cumulative analysis (n=13) of the frequency of CXCR5 and ICOS co-expression in TLR4^+^ (red) versus TLR4^-^ (grey) T cells. (J) Representative plots and cumulative analysis (n=13) of the frequency of ICOS and PD-1 co-expression TLR4^+^ (red) versus TLR4^-^ (grey) T cells. (K) Correlation between the frequency of TLR4^+^CXCR5^+^ T cells and TLR4^-^ CXCR5^+^ cells (n=13). (L) Correlation between the frequency of TLR4^+^PD1^+^ T cells and TLR4^-^PD1^+^ cells (n=13). ΔMFI was calculated to correct for the distinct autofluorescence of the TLR4^-^ and TLR4^+^ T cell populations. ΔMFI was calculated by subtracting the fluorescence intensity minus one (FMO) from median fluorescence intensity (MFI) for each given marker. D’Agostino & Pearson normality test was performed. p values ****p<0.0001, ***p<0.001, **p<0.01, *p<0.05 were determined by (B, C, D, E, F, H, I, J) Wilcoxon matched-pairs rank test; (G) Paired t-test; (K, L) Pearson Correlation.

These data indicate that TLR4^+^ T cells display Tfh-like features.

### TLR4^+^ T cells display migratory phenotype to inflamed tissues

TLR4^+^ T cell enrichment in synovial fluid (Fig. 1 G, H) cannot be fully explained by their CXCR5 expression. Therefore, we checked for the expression of chemokine receptors CCR2 and CCR6 that regulate T cell migration to inflamed tissues and whose ligands are abundantly present in arthritic synovium and have been implicated in the disease^43, 44^. Both CCR2 and CCR6 were upregulated by TLR4^+^ T cells (Fig 5 B-E). CCR2 and CCR6 are expressed by ∼100% and ∼30% of TLR4^+^ T cells, respectively (Fig. 5 B-E). While CCR2 guides a broad range of immune cells into sites of inflammation, CCR6 is associated with the recruitment of IL-17 producing T cells to inflamed joints^45^, suggesting an IL-17 inflammatory component to TLR4^+^ T cell synovial recruitment. To address this possibility, we checked whether TLR4^+^ T cells upregulate receptors for pro-inflammatory cytokines that are overexpressed in inflamed synovium (IL-1, IL-6 and IL-17) and which have been implicated in IL-17 production^45, 46^. IL-1R, whose engagement plays a critical in driving IL-17 mediated autoimmunity^46^, was selectively upregulated by TLR4^+^ T cells (Fig. 5 A, H, I). As expected from IL-6 pleiotropic role in immune responses, IL-6R was similarly expressed by both TLR4^+^ and TLR4^-^ T cell populations (Fig. 5 F, J, K). Finally, IL-17R, whose signaling reinforces IL-17 production and CD4^+^ T cells autoimmune profile^47^, was greatly enriched in TLR4^+^ T cells (Fig. 5 G, L, M). In addition, since TLR4^+^ T cells displayed a bigger cell size, we checked for the receptor of IL-2 alpha (IL-2R*α*), whose ligation drives T cell growth. IL-2R was increasingly expressed by TLR4^+^ (Fig. 5 N, O).

**Figure 5.**
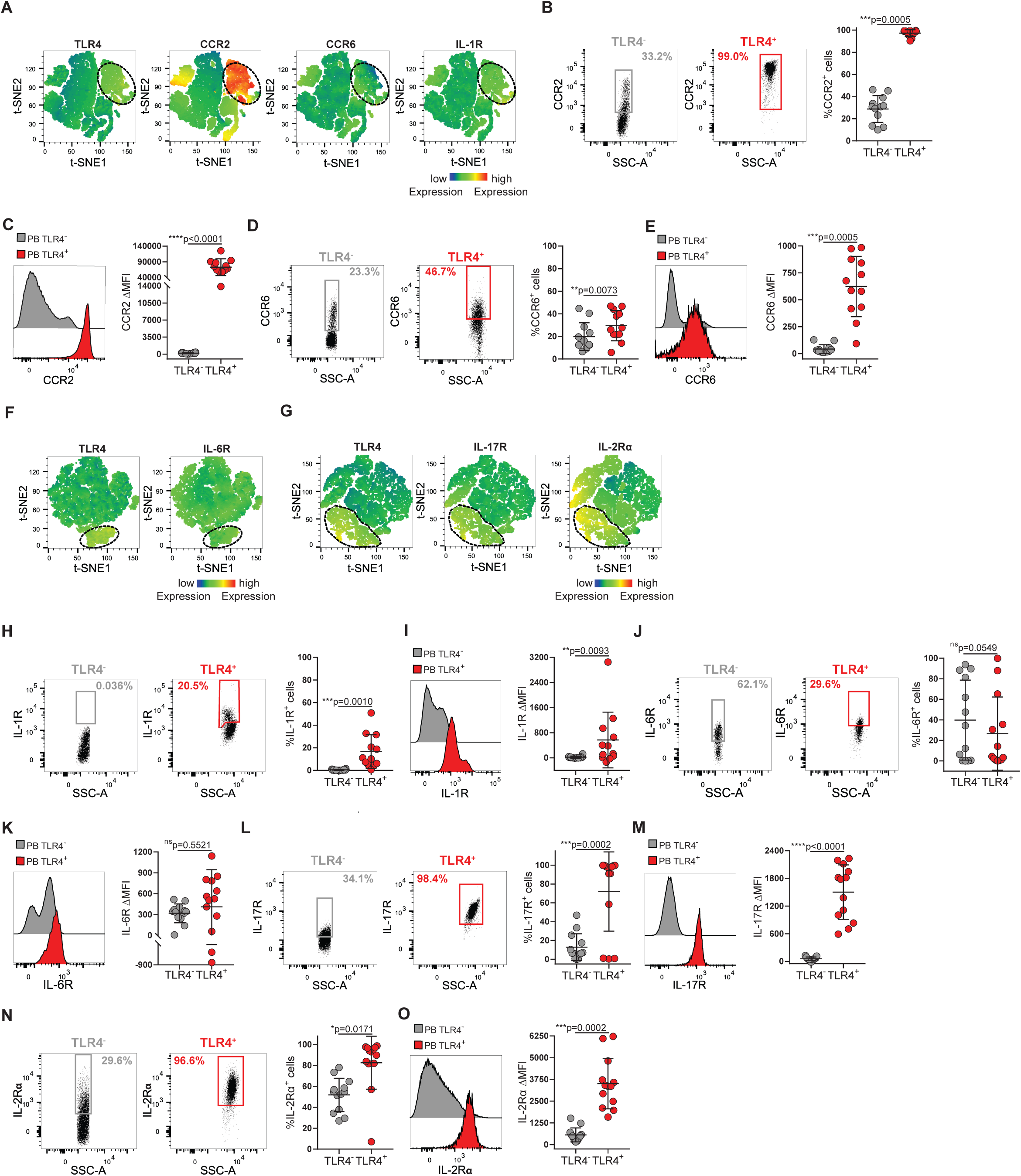
TLR4^+^ T cells display inflammatory chemokine and cytokine receptors. (A) t-SNE plots of peripheral blood total CD4^+^ T cells. Color indicates cell expression levels of labelled marker (TLR4, CCR2, CCR6, IL-1R). Circle demarks TLR4^+^ cells. (B, C) Representative plots and cumulative graph (n=12) of CCR2 frequency (B) and *Δ*MFI (C) in TLR4^+^ (red) and TLR4^-^ (grey) T cells. (D, E) Representative plots and cumulative graph (n=12) of CCR6 frequency (D) and *Δ*MFI (E) in TLR4^+^ (red) and TLR4^-^ (grey) T cells. (F-G) t-SNE plots of peripheral blood total CD4^+^ T cells. Color indicates cell expression levels of labelled marker. (F) TLR4, IL-6R. (G) TLR4, IL-17R and IL-2Rα. Circle demarks TLR4^+^ cells. (H, I) Representative plots and cumulative graph (n=12) of IL-1R frequency (H) and *Δ*MFI (I) in TLR4^+^ (red) and TLR4^-^ (grey) T cells. (J, K) Representative plots and cumulative graph (n=13) of IL-6R frequency (J) and *Δ*MFI (K) in TLR4^+^ (red) and TLR4^-^ (grey) T cells. (L, M) Representative plots and cumulative graph (n=13) of IL-17R frequency (L) and *Δ*MFI (M) in TLR4^+^ (red) and TLR4^-^ (grey) T cells. (N, O) Representative plots and cumulative graph (n=13) of IL-2Rα frequency (L) and *Δ*MFI (M) in TLR4^+^ (red) and TLR4^-^ (grey) T cells. ΔMFI was calculated to correct for the distinct autofluorescence of the TLR4^-^ and TLR4^+^ T cell populations. ΔMFI was calculated by subtracting the fluorescence intensity minus one (FMO) from median fluorescence intensity (MFI) for each given marker. D’Agostino & Pearson normality test was performed. p values ****p<0.0001, ***p<0.001, **p<0.01, *p<0.05 were determined by (B, E, H, I, J, L, N, O) Wilcoxon matched-pairs rank test; (C, D, K, M) Paired t-test.

Taken together TLR4^+^ T cells emerge as a Tfh-like cell population with a preferential tropism for inflamed tissues and increased capability to respond to IL-17 promoting stimuli. Curiously, TLR4^+^ T cells are not predisposed to respond to IL-6, which is involved in driving both Tfh cell differentiation^48^ and IL-17 production^45^, that is a current RA treatment target. Instead, TLR4^+^ T cells are predisposed to preferentially respond to IL-1 and IL-17. Due to the involvement of both IL-1R and IL-17R signaling in promoting chronic inflammatory IL-17 responses^46, 47^, selective increased expression of these receptors might ascribe a dysregulated IL-17 producing profile to TLR4^+^ T cells.

### TLR4 engagement reprograms TLR4^+^ T cell inflammatory profile

While in humans, the role of direct TLR4 engagement on CD4^+^ T cells remains largely unaddressed, in experimental autoimmune encephalitis, TLR4 engagement on CD4^+^ T cells has been reported to function as a co-receptor boosting T cell survival and proliferation without affecting the amount of the cytokines produced^31^. Whether or not direct TLR4 engagement on CD4^+^ T cells modulates or alters CD4^+^ T cell inflammatory profile has remained unanswered. To unveil the contribution of direct TLR4 engagement on T cell inflammatory profile, we FACS purified circulating CD4^+^ T cells from freshly obtained blood (Fig S1B; purity >99%) and stimulated them with highly purified TLR4 ligand LPS in the presence or absence of TCR and ICOS engagement. Since we had determined that TLR4^+^ T cells share features of Tfh-like cells and possess an IL-17 flavored pro-inflammatory phenotype, we looked at the antibody inducing cytokines which have been in RA pathology due to their role in promoting antibody production (IL-21^12^ and IL-10^60^), or in inducing joint tissue damage (IL-10^61^, IL-17^15, 19, 49, 50^ and TNF-*α*^59^). Circulating TLR4^+^ T cells produced IL-10, IL-21 and IL-17 in unstimulated conditions, supporting their ongoing activation state. In vitro, IL-21 production required TCR and ICOS stimulation and was completely non-responsive to LPS (Fig. 6 A, B). In contrast, LPS, in combination with TCR and ICOS stimulation, boosted IL-10, IL-17 and TNF-*α* production (Fig. 6 C-H). Moreover, LPS alone was sufficient to drive production of IL-10 and trended to increase IL-17 and TNF-*α* production, as well (Fig. 6 C, G).

**Figure 6.**
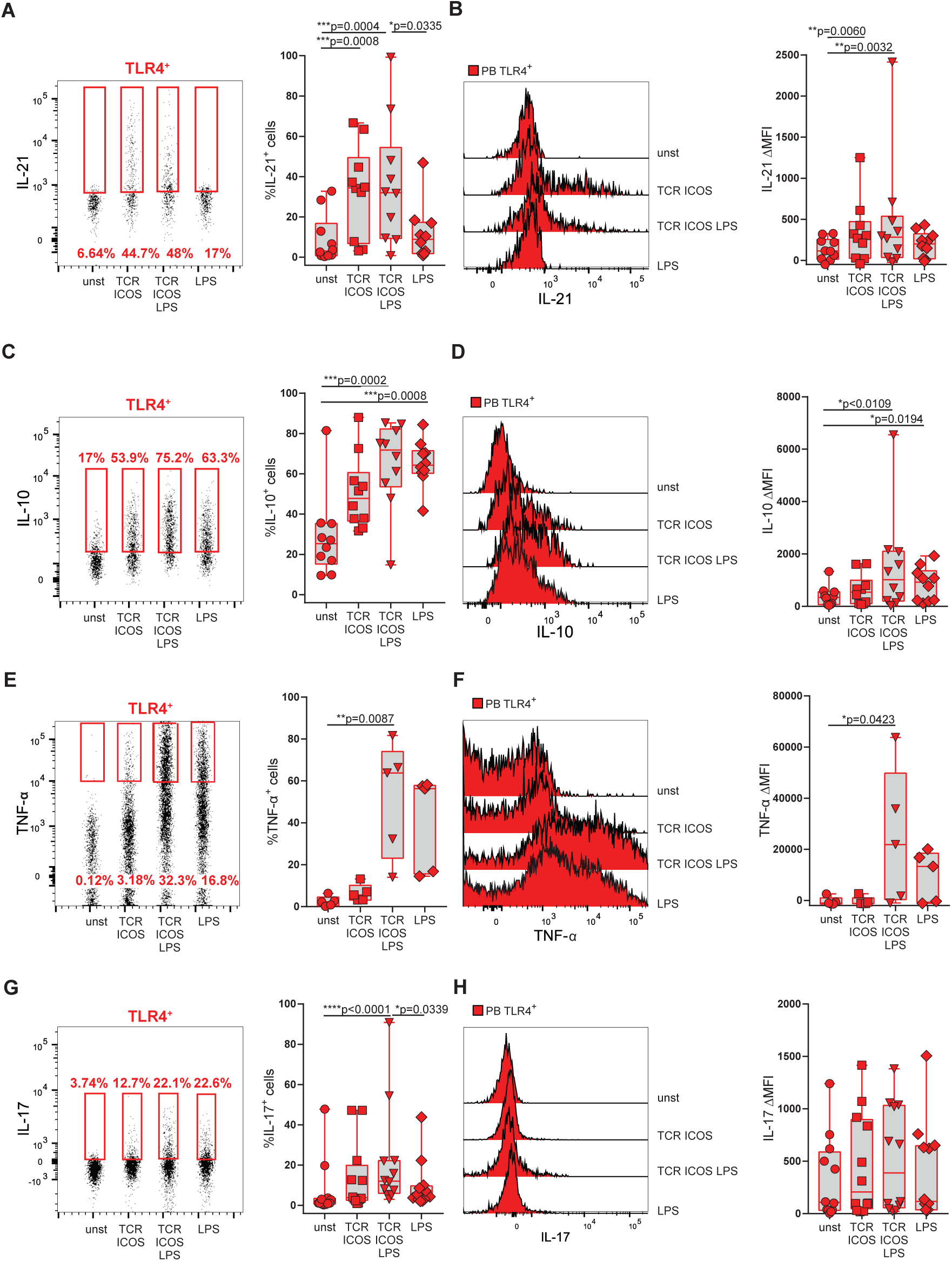
Direct recognition of LPS by TLR4^+^ T cells reprograms their cytokine program. FACS-purified CD3^high^CD4^high^ T cells from freshly obtained peripheral blood were cultured for 18 hours and stimulated with either *α*-CD3 and *α*-ICOS (TCR ICOS); *α*-CD3, *α*-ICOS and LPS (TCR ICOS LPS); LPS alone; or left unstimulated (unst). (A, B) Frequency (A) and *Δ*MFI (B) of IL-21 production by TLR4^+^ T cells (n=11). (C, D) Frequency (C) and *Δ*MFI (D) of IL-10 production by TLR4^+^ T cells (n=11). (E, F) Frequency (E) and *Δ*MFI (F) of TNF-*α* production by TLR4^+^ T cells (n=5). (G, H) Frequency (G) and *Δ*MFI (H) of IL-17 production by TLR4^+^ T cells (n=12). ΔMFI was calculated by subtracting the fluorescence intensity minus one (FMO) from median fluorescence intensity (MFI) for each given marker. D’Agostino & Pearson normality test was performed. p values ****p<0.0001, ***p<0.001, **p<0.01, *p<0.05 were determined by (A, B, C, D, E, F, G) Friedman test with posttest Dunn’s multiple comparisons; (H) RM one-way ANOVA with posttest Tukey’s multiple comparisons.

In sum, these data indicate that direct TLR4 stimulation goes beyond functioning as a coreceptor boosting TCR-driven response. While TCR and ICOS stimulation favors IL-21 production, LPS engagement shifts the inflammatory profile toward IL-17, TNF-*α* and of IL-10 production. Suggesting that TLR4 engagement by LPS might reprogram TLR4^+^ T cells from an IL-21 driven pro-antibody program to an inflammatory program fueling joint damage

### Direct recognition of TLR4 ligands present in synovial fluid drives IL-17 production, independently of antigen recognition

Increased expression of endogenous TLR4 ligands has been observed in the blood and synovial fluid of RA patients, with a role in arthritis being suggested in mice models^51–54^. Of all the proposed endogenous TLR4 ligands, tenascin-C is the one more thoroughly analyzed, including the molecular identification of its binding sites on TLR4^55^. Under physiological conditions, tenascin-C is tightly controlled, being virtually undetectable in healthy tissues, with transient re-expression occurring during tissue remodeling. Nonetheless, sustained tenascin-C accumulation occurs in a variety of chronic pathological conditions, including blood and synovial fluid of RA patients^26^. We quantified tenascin-C in synovial fluid of RA patients (Fig. 7 A-C). Synovial tenascin-C levels are independent of duration of DMARD treatment (Fig. 7 A), suggesting that tissue synovial deterioration persists despite of treatment. Moreover, TLR4^+^ T cells appear to be enriched in synovial fluids with higher tenascin-C levels (Fig. 7C), opening the possibility that tenascin-C might play a role in the enrichment of TLR4^+^ T cells in the synovial fluid. As we had observed that circulating TLR4^+^ T cells were producing IL-17 and IL-10 prior to in vitro restimulation (Fig. 6), we wondered whether this basal cytokine production was due to the ongoing engagement of TLR4. To address this possibility, we treated circulating TLR4^+^ T cells with either medium or with the TLR4 signaling inhibitor CLI-095. Blocking TLR4 signaling hampered both IL-17 and IL-10 production (Fig. 7 D-G). To further explore the role of direct TLR4 engagement by synovial components, we stimulated sorted CD3^high^CD4^high^ T cells with cell-depleted synovial fluid in the presence or absence of TLR4 signaling inhibitor. Stimulation with synovial fluid induced IL-17, IL-10 and TNF-*α*, but not IL-21, production. Increased IL-17, IL-10 and TNF-*α* production was mediated by direct TLR4 engagement, as it was abrogated by the addition of TLR4 specific signaling inhibitor CLI-095 (Fig. 7 H-J). In contrast to LPS (Fig. 6), direct TLR4 engagement by endogenous synovial ligands boosted IL-17, IL-10 and TNF-*α* production independently of TCR crosslinking. Reinforcing the view that endogenous TLR4 ligands and LPS elicit distinct inflammatory outcomes^26, 56, 57^. To scope the pathophysiological role that endogenous TLR4 ligands might exert on the inflammatory program of synovial TLR4^+^ T cells, we compared the cytokine profile of circulating and synovial TLR4^+^ T cells *ex vivo*. In this approach, freshly obtained and donor paired blood and synovial fluid mononuclear cells were immediately labelled for IL-17, IL-10, TNF-*α* and IL-21 (Fig. 7 L-O). Due to intrinsic differences in autofluorescence in blood and synovial fluid samples, FMOs were calculated independently for blood and for synovial fluid T cells. In all donors, ex vivo IL-17 production by TLR4^+^ T cells was higher in the synovium than in the blood (Fig. 7L). Curiously, IL-10 production is more prevalent in blood than in synovial TLR4^+^ T cells (Fig. 7M). Lastly, in our sampling we did not detect neither TNF-*α* nor IL-21 production by blood nor synovial TLR4^+^ T cells, *ex vivo*.

**Figure 7.**
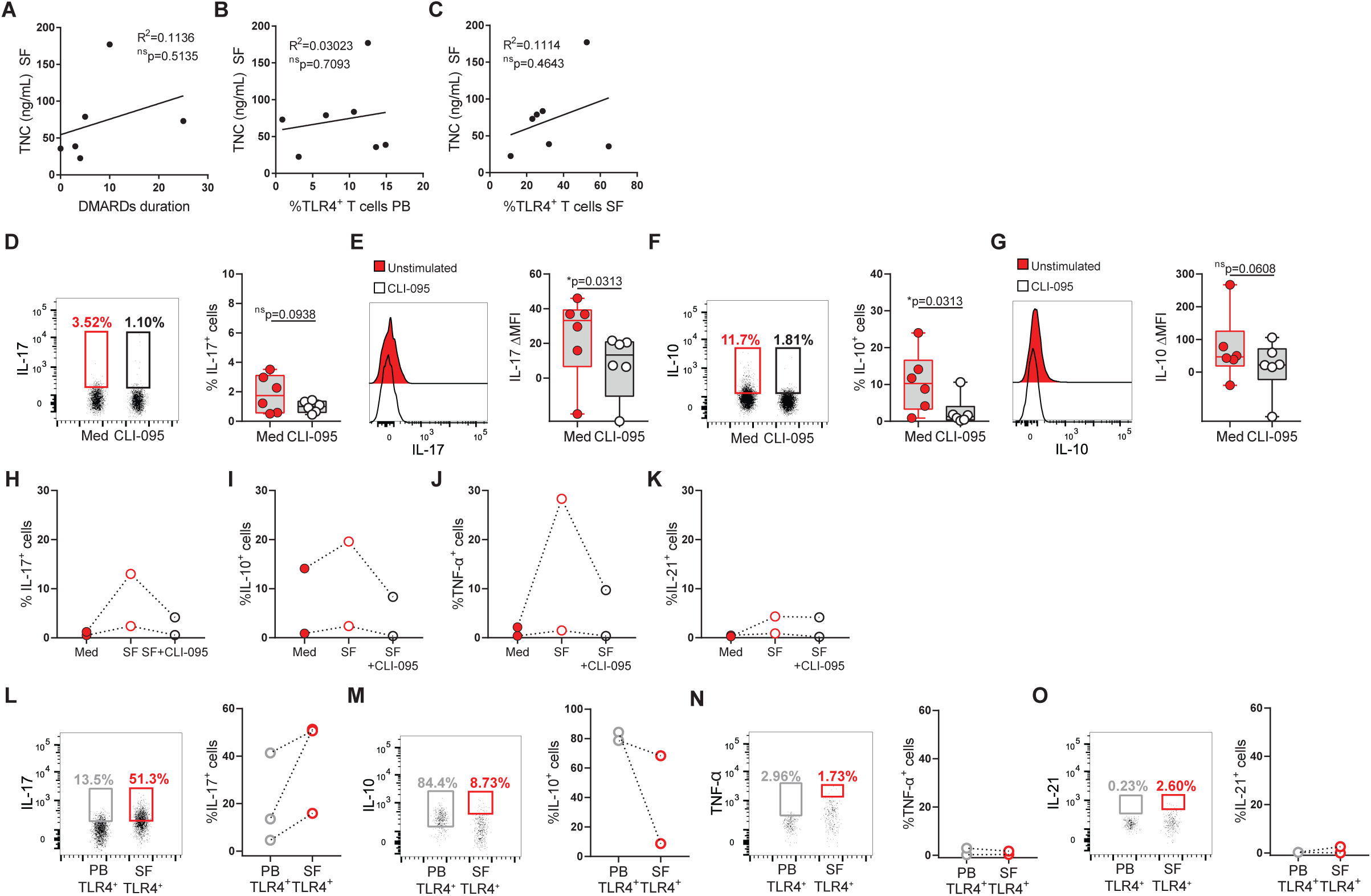
Direct recognition of TLR4 ligands present in synovial fluid drives IL-17 production, independently of antigen recognition. (A-C) Correlation between synovial fluid tenascin-C levels and DMARD duration (A, n=5), frequency of circulating (PB) TLR4^+^ T cells (B, n=7), and frequency of synovial fluid (SF) TLR4^+^ T cells (G, n=7). (D-G) FACS-purified CD3^high^CD4^high^ T cells from peripheral blood were cultured for 18 hours in the presence of medium (Med) or TLR4 signaling inhibitor (CLI-095). Frequency and *Δ*MFI of IL-17 (D, E) and IL-10 (F, G) production by TLR4^+^ T cells (n=6). (H-K) FACS-purified CD3^high^CD4^high^ T cells from peripheral blood were cultured for 18 hours in the presence of medium (Med), synovial fluid (SF) or TLR4 signaling inhibitor (CLI-095). Frequency IL-17 (H), IL-10 (I), TNF-*α* (J) and IL-21 (K) production by TLR4^+^ T cells (n=2). (L-O) Ex vivo production of IL-17, IL-10, TNF-*α* and IL-21 by TLR4+ T cells in n=3 freshly obtained peripheral blood (PB) and synovial fluid (SF) donor paired samples. ΔMFI was calculated by subtracting the fluorescence intensity minus one (FMO) from median fluorescence intensity (MFI) for each given marker. FMOs were calculated independently for blood and synovial fluid FACS analysis. Shapiro-Wilk normality test was performed because the n was too small for D’Agostino & Pearson normality testing. p values ****p<0.0001, ***p<0.001, **p<0.01, *p<0.05 were determined by (A, B, C) Pearson Correlation; (D, E, F) Wilcoxon matched-pairs rank test and (G) Paired t-test.

Altogether our results indicate that direct TLR4 engagement by endogenous ligands in synovial fluid favors the production of IL-17, IL-10 and TNF-*α*, but not IL-21. In contrast with LPS, endogenous synovial TLR4 ligands reprogram TLR4^+^ T cells inflammatory profile independently of TCR engagement. Lastly, cytokine production by synovial TLR4^+^ T cells suggest a major role for IL-17 in their pathogenic function.

## Discussion

RA is a chronic inflammatory disease where CD4^+^ T cells and joint tissue dysregulation synergize in propagating chronic inflammation and articular destruction. Treatment of RA remains challenging as identity of CD4^+^ T cell population driving RA and the mechanism by which joint microenvironment impinges dysregulated T cell activation remain elusive. Here, we identified a circulating TLR4^+^ T cell population which is enriched in synovial fluid of RA patients. TLR4^+^ T cells are uniquely attuned to respond distinctly to different contextual cues. They reconcile an unique ability to potentially promote systemic antibody production with an synovial-driven tissue damage program. Our results highlight the contribution of spatial compartmentalization, including homotypic cell:cell contacts, to T cell driven pathogenicity and the role of tissue environment in tailoring site-specific T cell responses.

Tfh-like cell populations have been described in several chronic inflammatory diseases including in rheumatoid arthritis^2^, lupus nephritis^58^ and systemic sclerosis^59^. In addition in RA, a population of IL-21 producing and antibody inducing peripheral helper T (Tph) cells has been identified^12^. Here we have identified a previously unknown Tfh-like population. TLR4^+^ T cells were enriched in Tfh cell markers, CXCR5 and PD-1^6–8^ and their frequency in circulation correlated with anti-CCP antibody levels. These sets of Tfh/Tph cells might indeed account for distinct cell populations or might represent the same cell population in different disease stages and/or response to treatment. Distinctly from previous reports^2, 12, 58, 59^, we analyzed freshly obtained blood and synovial fluid samples, rather than frozen ones. Fresh samples facilitate the identification of infrequent cell populations and the detection of certain markers and allow for a better detection of changes in cell size and shape.

Early descriptions of TLR4^+^ T had similarly reported an increase in cell size^60^. Likewise, *in vitro* and *in vivo* experiments show that IL-17 producing cells have a bigger size which has been associated with increased cytokine secretion in vitro^61^. As TLR4^+^ T cells FSC-A values were outside the conventional lymphocyte gate, we took care to exclude the occurrence of cell aggregates^62^. First, our doublet analysis (FSC-W vs FCS-A) into 2 distinct diagonals is suggestive of two cell populations rather than doublets. Second, confocal microscopy of FACS purified CD4^+^ T cells (∼99% purity) confirmed co-expression of TLR4 and CD3 exclusively by HLA-DR^+^FSC-A^high^ cells. TLR4 was expressed uniformly along the cell membrane, excluding the possibility of TLR4 acquisition through trogocytosis subsequent to prior interactions with APCs^63^. The ∼25% increase in cell size combined with membrane projections likely underpins to the 2-fold increase in FSC-A value detected by flow cytometry. Increase in cell size accompanied by expression of activation markers CD38 and HLA-DR further argues that TLR4^+^ T cells are indeed blasts.

HLA-DR are class II major histocompatibility molecules (MHC II) commonly present in APCs, where recognition of foreign-antigen bearing MHC by their cognate TCR on T cells drives antigen specific T cell activation^33, 64^. HLA-DR haplotypes constitute the strongest genetic association with RA^65^. So far, research addressing this genetic association has focused on identifying the immunodominant peptide presented by HLA-DR on APCs driving dysregulated T cell activation in RA patients. Even though, several citrullinated candidate peptides can be presented by HLA-DRB1^66^, the search for immunodominant T cell epitopes has so far revealed unfruitful. The observation that HLA-DR is expressed by activated T cells is longstanding^34, 35^, including a recent identification of a HLA-DR^+^ T cell subset in RA patients^67^. Nonetheless, the function of HLA-DR in T cells has remained enigmatic.

We unveiled for the first time a function for HLA-DR on T cells. By blocking MHCII:TCR interactions on FACS purified CD4^+^ T cells with an anti-HLA-DR antibody, we uncovered that homotypic T-T cell interactions through non-cognate HLA-DR:TCR contacts regulate TLR4 surface expression on T cells. Homotypic T:T cell interactions have been described to be established through adhesion molecules^68^ and to support T cell differentiation and enhancement of the immune response^69^. We have broadened the range of molecules involved in homotypic T:T cell interactions by identifying that they can be additionally established through non-cognate HLA-DR:TCR interactions. Non-cognate HLA-DR:TCR interactions between APCs and T cells are known to alter T cell genetic profile^40, 41^. Thus, it is possible that homotypic T-T cell interactions through non-cognate HLA-DR:TCR contacts might drive TLR4 gene expression. Another possibility is that these homotypic HLA-DR:TCR interactions stabilize TLR4 expression at the T cell plasma membrane. Further studies will be needed to dissect the mechanism by which homotypic HLA-DR:TCR interactions regulate TLR4 expression on T cells. It is possible that these homotypic HLA-DR:TCR interactions occur more frequently in the densely packed joint environment, where TLR4^+^ T cells are enriched. Suggesting the enticing possibility that HLA-DR mediated homotypic interactions might sensitize for joint microenvironment recognition and for contextually driven shift of their pathological program.

TLR4 is a relatively promiscuous immune sensor that recognizes both microbial and endogenous ligands. In ankylosing spondylitis patients, CD28^-^TLR4^+^ T cells with a senescent phenotype were found to be present in the blood but practically absent from affected joints^39^. This is in stark contrast with the TLR4^+^ Tfh-like cell population reported here; TLR4^+^ T cells were expanded in synovial fluid, and even though they were enriched for PD-1 they did not exhibit signs of wither exhaustion or senescence, as illustrated by their highly proliferative status and increased ability to produce cytokines in response to stimulation. In addition to CXCR5, TLR4^+^ T cells also expressed the chemokine receptors CCR2 and CCR6 indicating a preferential recruitment to inflamed tissues, which might account for their enrichment in the affected joints. Interestingly, TLR4 signaling has been reported to augment cell migration and invasiveness^70, 71^, opening the possibility that direct TLR4 engagement could propel T cell invasiveness into affected joint.

In mice models of autoimmune diseases TLR4 signaling in CD4^+^ T cells has been reported to function both as disease facilitator^31^ and protector^30^. Nonetheless, a role for direct TLR4 engagement in T cell cytokine profile and function had not been reported so far. Our data show that while TCR engagement favors production of antibody inducing cytokine IL-21, TLR4 engagement by either LPS or synovial fluid components ensues IL-17, IL-10 and TNF-*α* production, cytokine whose role in RA has been ascribed to promoting joint damage^15, 19, 49, 50, 59, 61^. Even though IL-10 is often labeled as an anti-inflammatory cytokine, it is well established that IL-10 has both immunosuppressive and stimulatory effects, including cytotoxic activity against tumors^72^. In RA, IL-10 has been reported to drive inflammatory arthritis and joint destruction^73^. The existence of an antibody-independent pathogenic function for TLR4^+^ T cells would explain why this population is also present in seronegative RA patients.

Curiously, while TLR4 engagement by LPS functions as a costimulatory signal boosting TCR signaling, TLR4 ligation by endogenous TLR4 ligands fuels TLR4^+^ T cell inflammatory program independently of cognate antigen recognition. Distinct ligands ensuing different TLR4 responses is likely due to the fact that TLR4 has multiple binding sites^55^. In fact, TLR4 ligation by endogenous ligands Tenascin-C and fibronectin is not blocked by an LPS mimetic, which blocks TLR4 activation by competing with LPS for TLR4/MD-2 binding^26, 74^. In addition, gene expression profiles induced by hyaluronan and tenascin-C are significantly different from that induced by LPS^26, 56, 57^. Even though, we cannot formally exclude that other components present in the synovial fluid might affect T cell function, blocking of TLR4 in the presence of synovial fluid completely abrogated IL-17, IL-10 and TFN-*α* production. Thus, we can conclude that the production of IL-17, IL-10 and TNF-*α* induced by synovial fluid is specifically mediated by TLR4 on T cells. It is likely that these TLR4 sponsored effects are mediated by the combined action of several endogenous TLR4 ligands present in the joints.

Importantly, *ex vivo* freshly analyzed synovial TLR4^+^ T cells seemed to be skewed toward IL-17 production. When compared to in vitro stimulation with cell-depleted synovial fluid, synovial TLR4^+^ T cells seemed to be poised to produce more IL-17, less IL-10 and no TNF-*α*. These differences might be due to the fact that to release cells from synovial fluid, it is necessary to degrade it enzymatically. Hyaluronidase digestion could give rise to additional TLR4 ligands that could be more adept at inducing IL-10 and TNF-*α* in in vitro restimulation assays. In particular, different molecular weight hyaluronic acid fragments are known to elicit distinct inflammatory profiles^75^. It is possible that in vivo, IL-17 is the main cytokine induced by direct engagement of TLR4 on synovial T cells, where it might play a prominent role in mediating bone erosions and cartilage damage^76, 77^.

Our study recruited a considerable RA patient cohort of 103 patients. Nonetheless, there are some limitations to our study. We could only obtain a relatively modest number of synovial fluid samples. This was due to the fact that we only used freshly obtained synovial fluid whose access to was seriously hindered during this last year COVID-19 pandemic imposed serious restrictions on hospital access to chronic patients. Another limitation was that most of the patients recruited were either in remission or presented controlled disease and thus had very low DAS scores, which made difficult to correlate the frequency of TLR4^+^ T cells with disease activity. Despite these limitations, TLR4^+^ T cells in the blood and synovial fluids correlated well indicating that the blood can be used to probe TLR4^+^ T cell synovial enrichment. In addition, our functional assays were robust identifying a causal relationship linking TLR4^+^ T cells selective recognition of joint tissue environment to the type immune profile ensued. Further studies will be needed to address the impact of TLR4^+^ T cells in joint damage.

Deciphering which CD4^+^ T cells are relevant to the disease process and they interplay with the joint microenvironment is a critical hurdle to our understanding of RA. Here we propose a mechanism by which the joint tissue microenvironment might reset on TLR4^+^ T cells pathological function. Outside the joints, TLR4^+^ Tfh-like cells will be activated predominantly through the TCR leading to the production of IL-21, which favors antibody production and will likely contribute to anti-CCP antibody titers (Fig. 8). Within in the affected joints homotypic T:T cell interactions mediated through non-cognate HLA-DR:TCR coupling supports TLR4 surface expression. In turn, direct sensing of joint damage patterns by TLR4^+^ T cells reprograms them towards an IL-17 pathological program that drives and sustains cartilage damage and bone erosions (Fig.8). This two-prong mechanism could highlight several attractive therapeutic targets both at the systemic level and in the affected tissues. In addition, circulating TLR4^+^ T cells could constitute a good biomarker to predict flares and possibly which patients are more likely to develop cartilage damage and joint erosions.

**Figure 8.**
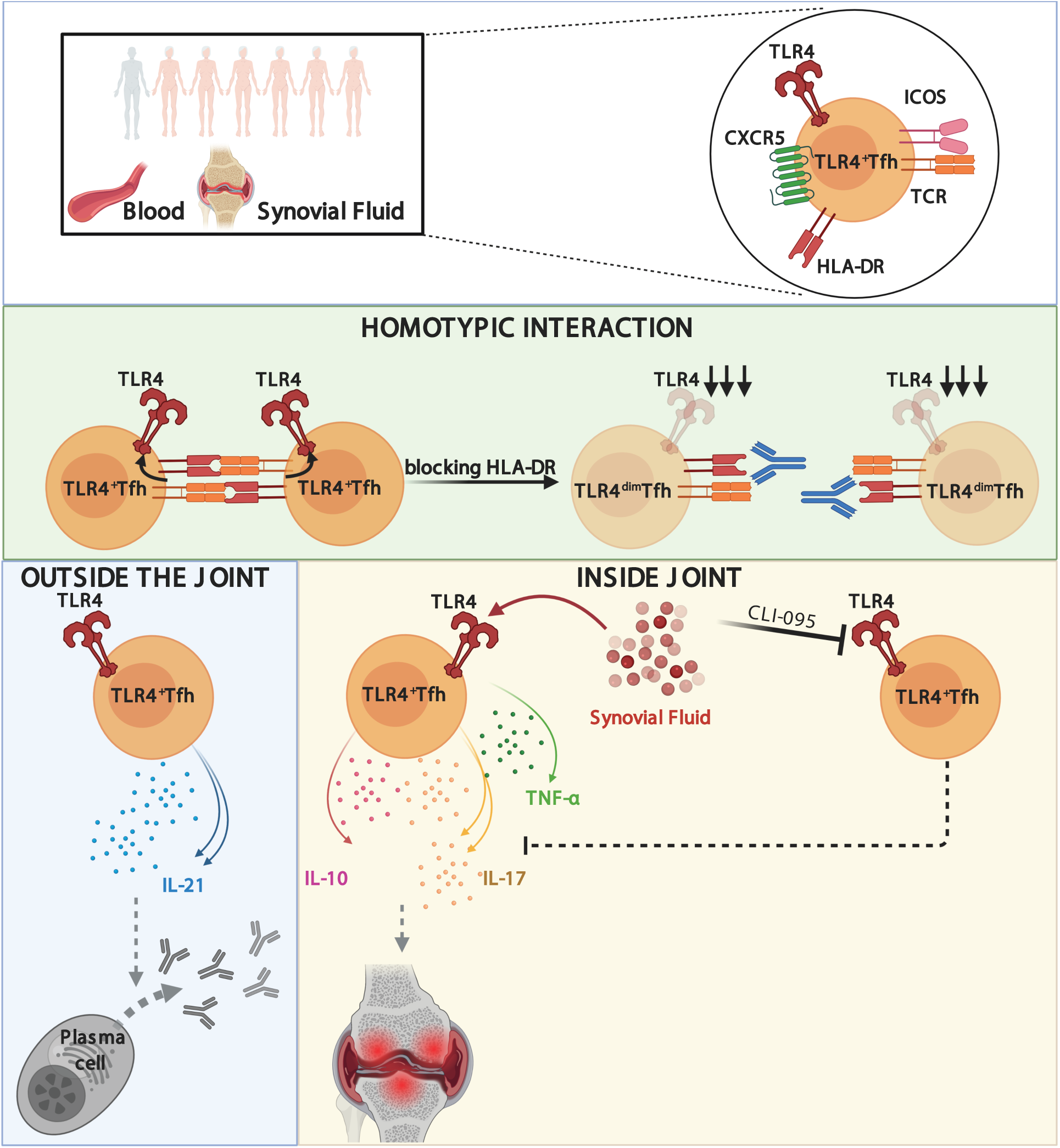
Schematic representation of proposed mechanism and role of TLR4^+^ T cells in RA. From top to bottom: a population of TLR4^+^ Tfh-like T cells expanded in synovial fluid of RA patients was characterized in a cohort of more than 100 RA patients. We uncovered a function for HLA-DR expression by T cells, as homotypic T:T cell contacts established through HLA-DR:TCR were found to be required for TLR4 expression by T cells. TLR4^+^ T cells display a two-pronged mechanism uniquely poised to tailor their pathogenic phenotype in response to contextual cues. Outside the joints, TCR driven IL-21 production favors antibody production, which likely contributes to anti-CCP antibody titers. Within in the affected joints direct sensing of joint damage patterns by TLR4^+^ T cells reprograms them towards an IL-17 pathological program known to drive and sustain cartilage damage and bone erosions.

## Acknowledgements

We thank Cláudia Andrade for technical support and Juliana Gonçalves for testing samples for SARS-CoV-2 exposure. We are extremely grateful to all the participants of the study and to the whole rheumatology department at Hospital Egas Moniz that made this study possible. This work was supported by FCT grant (PTDC/MEC-REU/29520/2017 to H.S.) and by iNOVA4Health (Grant UID/Multi/04462 to HS). HS is supported by Fundação para a Ciência e Tecnologia (FCT) under the FCT Investigator Program (IF/01722/2013) and by H2020-WIDESPREAD-01-2016-2017-TeamingPhase2 - 739572 - THE DISCOVERIES CTR, DAS and RCT were supported by FCT through PD/BD/137409/2018 and UID/Multi/04462, respectively.

## Author contributions

DAS and RCT designed and performed experiments. DAS analyzed the data. RT collected clinical data. AN, IS, SF, MC, NPG, RT, MJG recruited patients and provided blood and synovial recruited patients, provided blood and synovial fluid samples. ABS, CL, MM, MHL, PA, SM, TC, WC recruited patients and provided blood samples. FPS and AFM recruited patients, provided blood samples, and discussed clinical data. JCB advised, analyzed and interpreted clinical data. HS conceived the project, designed and performed experiments, supervised the project, analyzed the data and wrote the manuscript. All authors discussed the results and commented on the manuscript.

## METHODS

### Human samples

The Ethics Committee of NOVA Medical School and of Hospital Egas Moniz approved this study. Informed consent was obtained from RA patients that fulfilled ACR 2010 classification criteria. Rheumatoid factor status, C-reactive protein level, erythrocyte sedimentation rate and medication usage were obtained by review of medical records. Anti-CCP antibody titers were determined at the time of blood draw using a commercial assay anti-CCP ELISA (IgG) from EUROIMMUN with a positive result defined as >5RU/mL. Number of swollen and/or tender joints was measured by attending clinician on the day of sample acquisition. Treatments are categorized in: non-steroid anti-inflammatory (NSAID), corticosteroids, disease modifying antirheumatic drugs (DMARDs) and biological DMARDs (dDMARDs). Blood was drawn by venipuncture into Lithium-Heparin containing cell preparation tubes (BD, Vacutainer). Synovial fluid was collected only when excess material from patients undergoing diagnostic or therapeutic arthrocentesis. Demographic and clinical data for all the patients enrolled in this study are listed in Table S1.

### Peripheral blood and synovial fluid cell isolation

Blood samples and synovial fluid were processed within 2 hours of collection and freshly analyzed. Peripheral blood and synovial mononuclear cells were isolated by density gradient centrifugation (Biocoll, Merck Millipore) or following enzymatic digestion with hyaluronidase (10µL/mL; 30min at 37°C), respectively. Plasma and cell-depleted synovial fluid were frozen until further use.

### Antibodies and flow cytometry

For flow cytometry analysis peripheral blood cells were stained with antibodies listed in Table S2. For cell viability, Fixable Viability Dye (eBioscience) or Calcein Violet-AM (Biolegend) were used. When described, cells were cultured overnight with 10 μg/mL of anti-HLA-DR antibody (L243). For intracellular staining cells were treated with Transcriptional Factor Fixation/Permeabilization kit (ebioscience). FACS acquisition was performed in a BD FACSCanto II instrument (BD Biosciences) and further analyzed with FlowJo v10.7.1 software.

### Cell sorting and intracellular cytokine staining

For flow cytometry cell sorting, cells were stained with anti-CD4 (RPA-T4) and anti-CD3 (SK7) antibodies (BioLegend) or with anti-CD4 (RPA-T4), anti-CD3 (SK7), anti-HLA-DR (L243). Gating strategies are depicted in Fig. S1 A, B. Sorted populations cell purity was routinely >98% (Fig. S1C). For intracellular cytokines assays sorted CD3^high^CD4^high^, rested for at least 3h, were stimulated with 5 μg/mL of anti-CD3 (UCHT1, BioLegend) and 2 μg/mL of anti-ICOS (C398.4A, BioLegend), crosslinked with 5 μg/mL anti-mouse IgG1 (BioLegend) plus 10 μg/mL anti-hamster IgG (Thermo Fisher Scientific) at 37°C in the presence of Brefeldin-A (Life Technologies) for 14 h. Cells were fixed in paraformaldehyde 1% (Sigma-Aldrich) and permeabilized with saponin (Carl Roth). Antibodies used are listed in Table S2. When indicated 1.7 μg/mL LPS (Sigma-Aldrich) or cell-depleted synovial fluid (SF) was added. For TLR4 blocking, CLI-095 (InvivoGen) was added at 10 μg/mL 1h before stimulation. Cell sorting was performed in a BD FACSAria III instrument (BD Biosciences).

### Imaging, image processing, and quantification

FACS-purified CD3^high^CD4^high^HLA-DR^+^ cells were immediately plated onto poly-L-lysine–coated coverslips, fixed in 4% paraformaldehyde for 15 min at room temperature, incubated with blocking buffer (PBS BSA 1%) and immunostained as previously described^33, 78^. Antibodies used for immunofluorescence staining are described in Table S2. Confocal images were obtained using a Zeiss LSM 710 confocal microscope (Carl Zeiss) over a 63x objective. Z stack optical sections were acquired at 0.2 μm depth increments, and both green and red laser excitation were intercalated to minimize crosstalk between the acquired fluorescence channels. 3D image deconvolution was performed using Huygens Essential 19.10, and 2D images were generated from a maximum intensity projection over a 3D volume cut of 0.4-μm depth centered on the cell medium plane using Imaris. For quantification of cell size and roundness, confocal images were acquired at 2-μm increments in the z-axis.

### Flow Cytometry Data analysis

Flow cytometry data was analyzed using FlowJo and pluggins DownSample and FlowAI. The flow cytometry data was compensated at the time of acquisition with UltraComp eBeads (Thermo Fisher). As controls unstained and fluorescence minus one (FMO) conditions were included. The data collected in .fcs files was analyzed so that all abnormal events would be excluded by using FlowAI (Gianni Monaco et al. flowAI: automatic and interactive anomaly discerning tools for flow cytometry data. Bioinformatics 2016, 1-8 https://doi.org/10.1093/bioinformatics/btw191). Then, by using the gating strategies mentioned in the figures, dead cells and doublets were excluded. Whenever mentioned *Δ*MFI was calculated by subtracting the Fluorescence Minus One (FMO) FMO from MFI for any given fluorophore being analyzed. t-SNE maps were generated by pooling patients. Every heatmap represents differential marker expression between TLR4^+^ cells (dashed gate) and remaining CD4^+^ T cell populations. To maintain the consistency of the events from each condition and also to reduce the number of events fed into t-SNE algorithm, DownSample was used and files were concatenated in a way that all conditions/donors could be represented in the same plot.

### Statistical analysis

Results are presented as medians. GraphPad Prism v8.4.2 software was used for statistical analysis. To test the normality of the data, D’Agostino & Pearson normality test was used. In two groups comparison: for paired data, Paired t-test or Wilcoxon matched-pairs signed rank test was used; for unpaired data, Mann-Whitney test was used. For multiple groups comparison: for unpaired data Krustall-Wallis test with posttest Dunn’s multiple comparisons; for paired data, RM one-way ANOVA with posttest Turkey’s multiple comparisons or Friedman test with posttest Dunn’s multiple comparisons were used as indicated. For correlations Pearson or Spearman was used as described. Results were considered significant at *p < 0.05, **p < 0.01, ***p < 0.001, ****p <0.0001.

## Supplemental information

**Figure S1.**
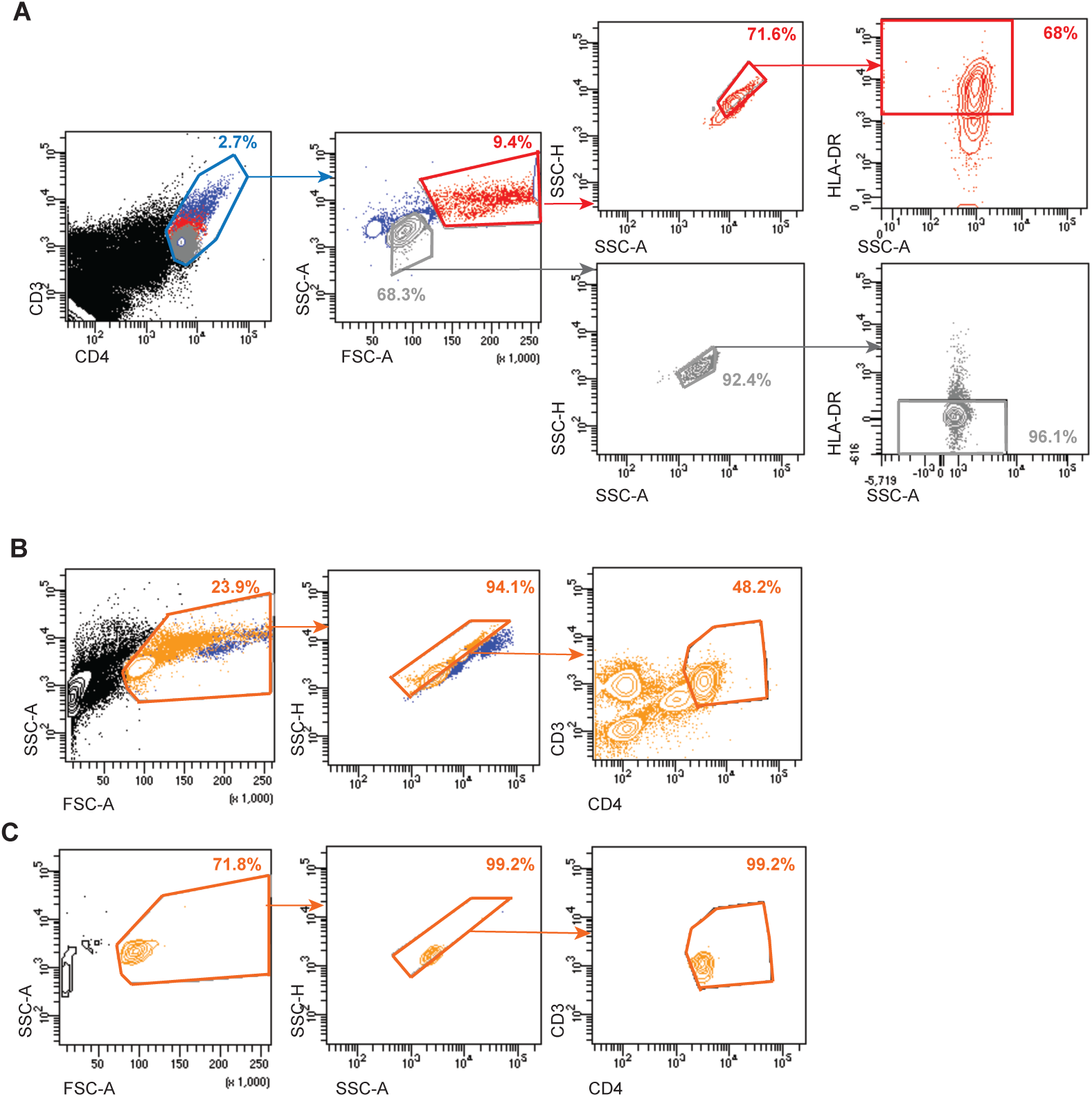
FACS-purification strategy and sorted cell population purity. (A)Flow cytometric sorting strategy for the purification of CD3^high^CD4^high^HLA-DR^+^ and CD3^high^CD4^high^HLA-DR^-^ T cells. (B) Flow cytometric sorting strategy for the purification of CD3^high^CD4^high^ T cells. (C) Purity of sorted CD3^high^CD4^high^ T cells.

**Table 1.**
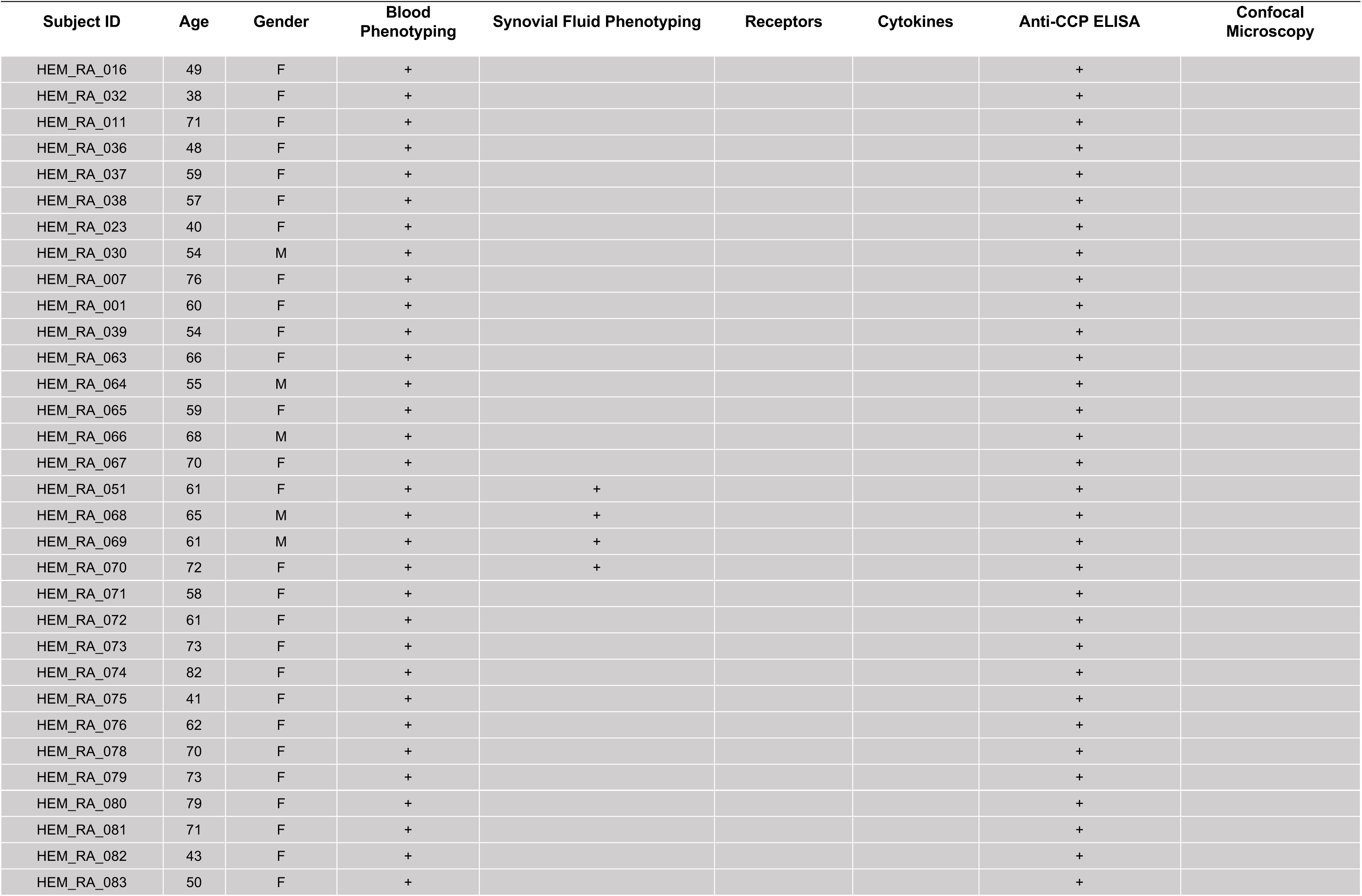
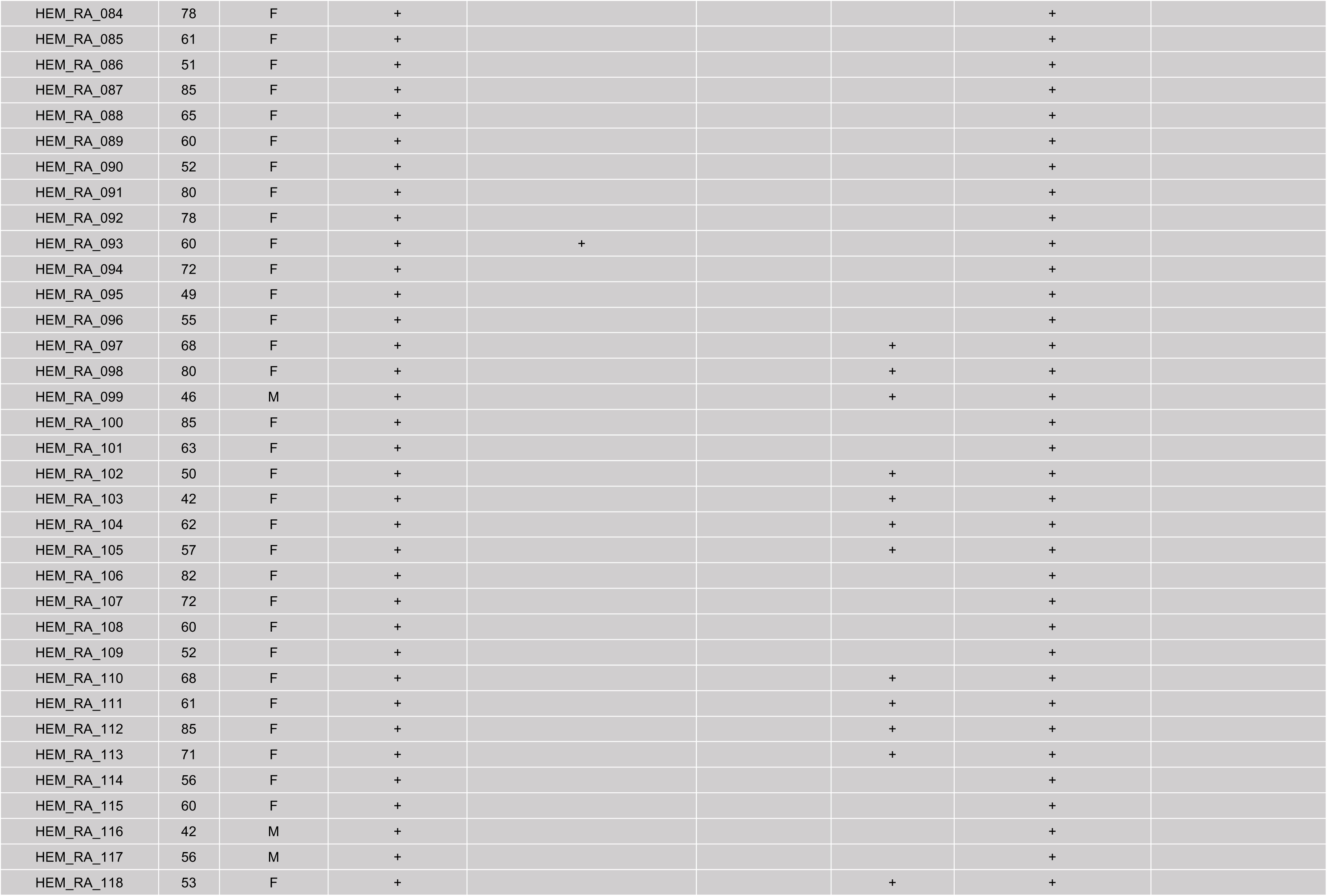
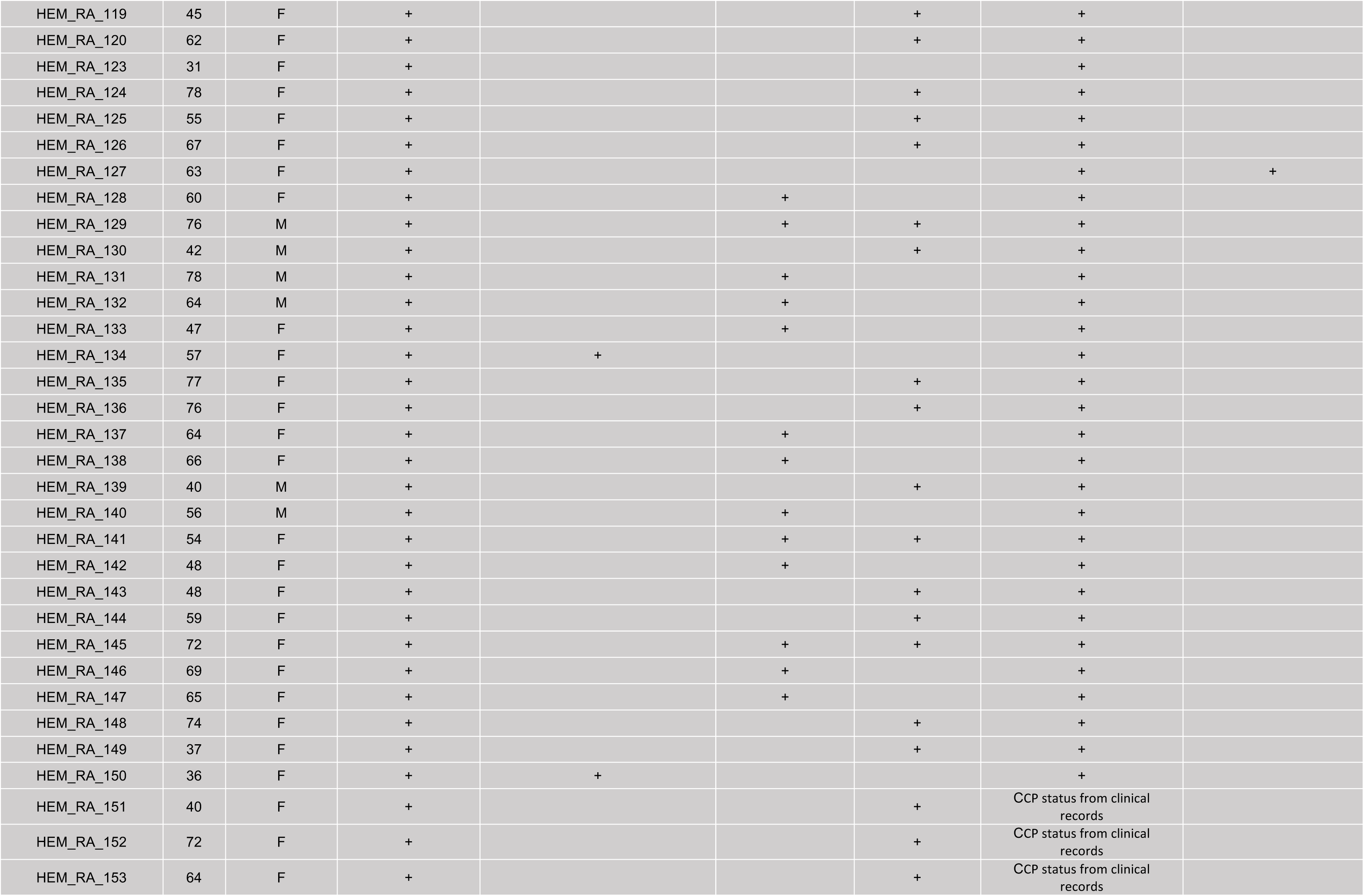
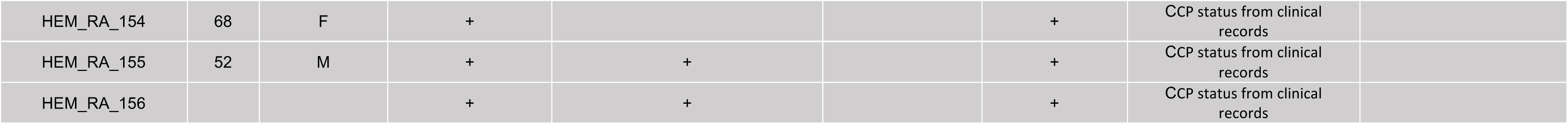
Donors.

**Table 2.**
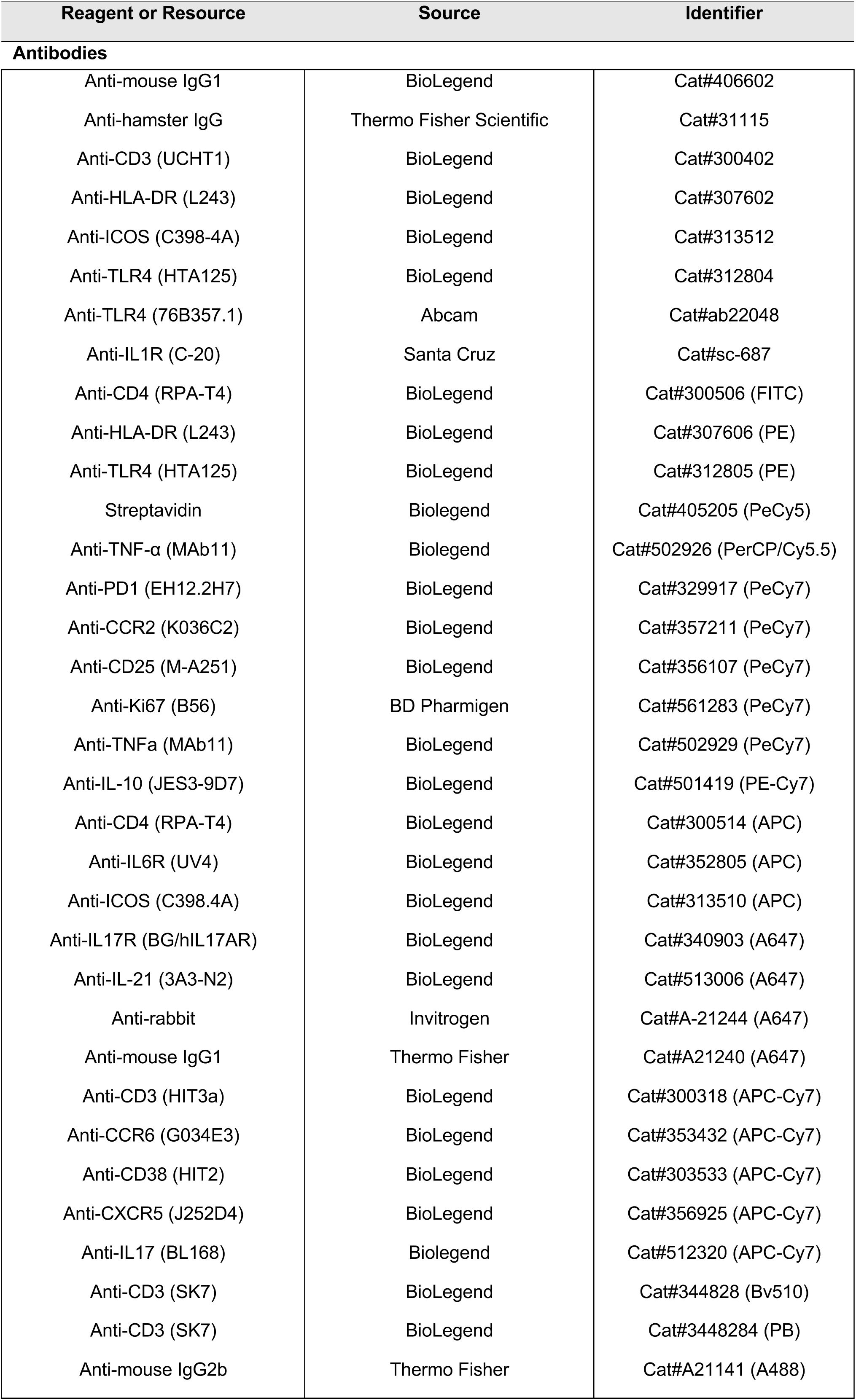
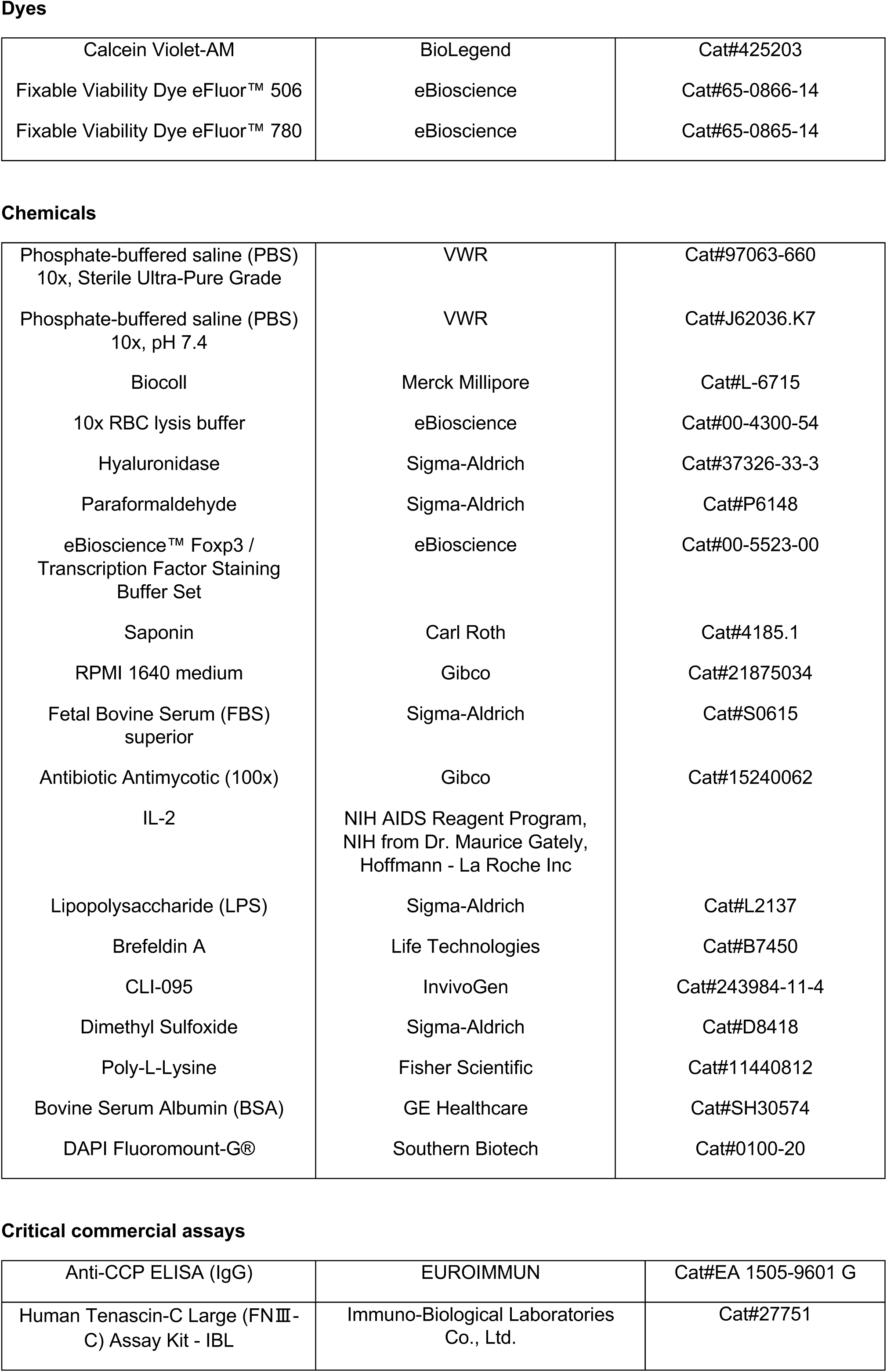
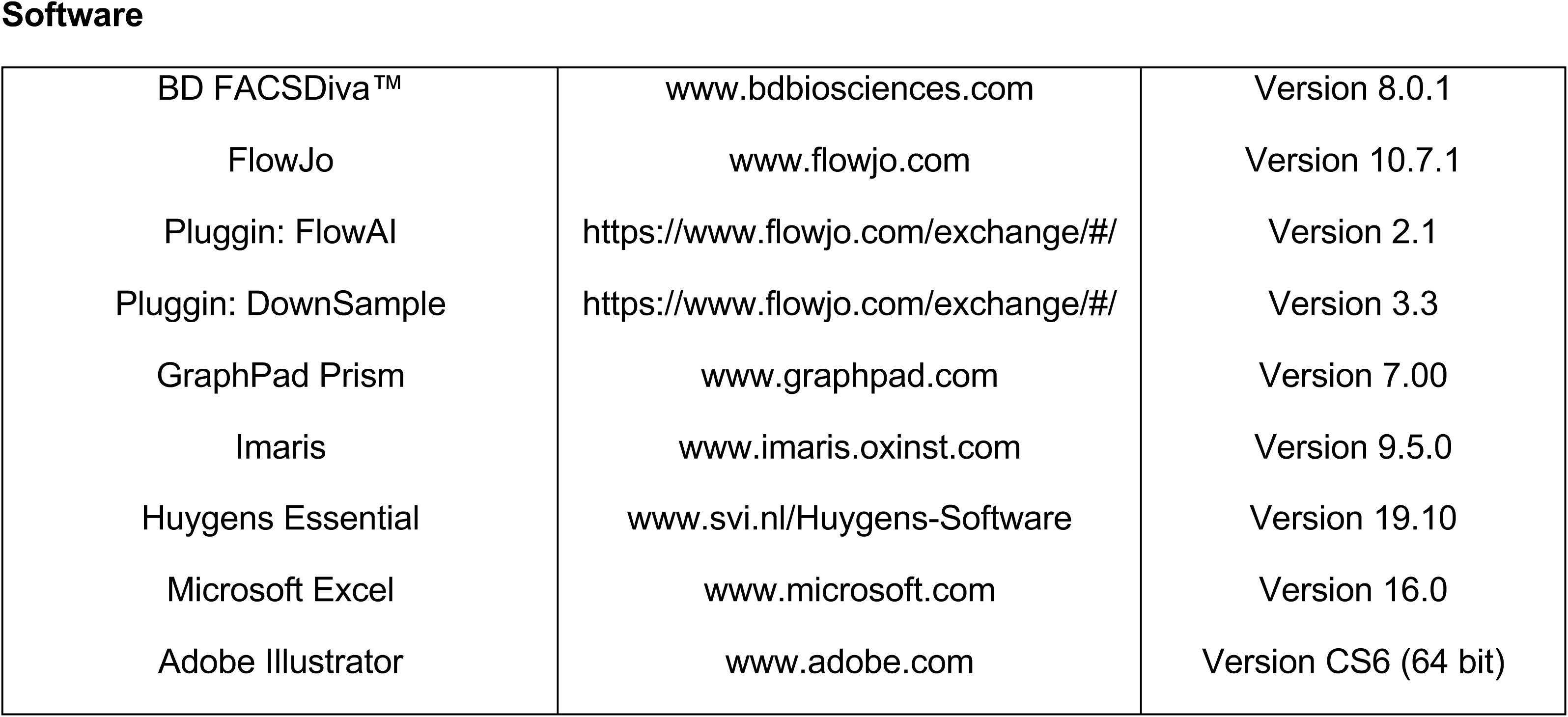
Reagents.

| Reagent or Resource | Source | Identifier |
| --- | --- | --- |
| **Antibodies** |  |  |
| Anti-mouse IgG1 | BioLegend | Cat#406602 |
| Anti-hamster IgG | Thermo Fisher Scientific | Cat#31115 |
| Anti-CD3 (UCHT1) | BioLegend | Cat#300402 |
| Anti-HLA-DR (L243) | BioLegend | Cat#307602 |
| Anti-ICOS (C398-4A) | BioLegend | Cat#313512 |
| Anti-TLR4 (HTA125) | BioLegend | Cat#312804 |
| Anti-TLR4 (76B357.1) | Abcam | Cat#ab22048 |
| Anti-IL1R (C-20) | Santa Cruz | Cat#sc-687 |
| Anti-CD4 (RPA-T4) | BioLegend | Cat#300506 (FITC) |
| Anti-HLA-DR (L243) | BioLegend | Cat#307606 (PE) |
| Anti-TLR4 (HTA125) | BioLegend | Cat#312805 (PE) |
| Streptavidin | Biolegend | Cat#405205 (PeCy5) |
| Anti-TNF-α (MAb11) | Biolegend | Cat#502926 (PerCP/Cy5.5) |
| Anti-PD1 (EH12.2H7) | BioLegend | Cat#329917 (PeCy7) |
| Anti-CCR2 (K036C2) | BioLegend | Cat#357211 (PeCy7) |
| Anti-CD25 (M-A251) | BioLegend | Cat#356107 (PeCy7) |
| Anti-Ki67 (B56) | BD Pharmigen | Cat#561283 (PeCy7) |
| Anti-TNFA (MAb11) | BioLegend | Cat#502929 (PeCy7) |
| Anti-IL-10 (JES3-9D7) | BioLegend | Cat#501419 (PE-Cy7) |
| Anti-CD4 (RPA-T4) | BioLegend | Cat#300514 (APC) |
| Anti-IL6R (UV4) | BioLegend | Cat#352805 (APC) |
| Anti-ICOS (C398.4A) | BioLegend | Cat#313510 (APC) |
| Anti-IL17R (BG/hIL17AR) | BioLegend | Cat#340903 (A647) |
| Anti-IL-21 (3A3-N2) | BioLegend | Cat#513006 (A647) |
| Anti-rabbit | Invitrogen | Cat#A-21244 (A647) |
| Anti-mouse IgG1 | Thermo Fisher | Cat#A21240 (A647) |
| Anti-CD3 (HIT3a) | BioLegend | Cat#300318 (APC-Cy7) |
| Anti-CCR6 (G034E3) | BioLegend | Cat#353432 (APC-Cy7) |
| Anti-CD38 (HIT2) | BioLegend | Cat#303533 (APC-Cy7) |
| Anti-CXCR5 (J252D4) | BioLegend | Cat#356925 (APC-Cy7) |
| Anti-IL17 (BL168) | Biolegend | Cat#512320 (APC-Cy7) |
| Anti-CD3 (SK7) | BioLegend | Cat#344828 (Bv510) |
| Anti-CD3 (SK7) | BioLegend | Cat#3448284 (PB) |
| Anti-mouse IgG2b | Thermo Fisher | Cat#A21141 (A488) |
| **Dyes** |  |  |
| Calcein Violet-AM | BioLegend | Cat#425203 |
| Fixable Viability Dye eFluor™ 506 | eBioscience | Cat#65-0866-14 |
| Fixable Viability Dye eFluor™ 780 | eBioscience | Cat#65-0865-14 |
| **Chemicals** |  |  |
| Phosphate-buffered saline (PBS)<br>10x, Sterile Ultra-Pure Grade | VWR | Cat#97063-660 |
| Phosphate-buffered saline (PBS)<br>10x, pH 7.4 | VWR | Cat#J62036.K7 |
| Biocoll | Merck Millipore | Cat#L-6715 |
| 10x RBC lysis buffer | eBioscience | Cat#00-4300-54 |
| Hyaluronidase | Sigma-Aldrich | Cat#37326-33-3 |
| Paraformaldehyde | Sigma-Aldrich | Cat#P6148 |
| eBioscience™ Foxp3 /<br>Transcription Factor Staining<br>Buffer Set | eBioscience | Cat#00-5523-00 |
| Saponin | Carl Roth | Cat#4185.1 |
| RPMI 1640 medium | Gibco | Cat#21875034 |
| Fetal Bovine Serum (FBS)<br>superior | Sigma-Aldrich | Cat#S0615 |
| Antibiotic Antimycotic (100x) | Gibco | Cat#15240062 |
| IL-2 | NIH AIDS Reagent Program,<br>NIH from Dr. Maurice Gately,<br>Hoffmann - La Roche Inc |  |
| Lipopolysaccharide (LPS) | Sigma-Aldrich | Cat#L2137 |
| Brefeldin A | Life Technologies | Cat#B7450 |
| CLI-095 | InvivoGen | Cat#243984-11-4 |
| Dimethyl Sulfoxide | Sigma-Aldrich | Cat#D8418 |
| Poly-L-Lysine | Fisher Scientific | Cat#11440812 |
| Bovine Serum Albumin (BSA) | GE Healthcare | Cat#SH30574 |
| DAPI Fluoromount-G® | Southern Biotech | Cat#0100-20 |
| **Critical commercial assays** |  |  |
| Anti-CCP ELISA (IgG) | EUROIMMUN | Cat#EA 1505-9601 G |
| Human Tenascin-C Large (FNIII-C)<br>Assay Kit - IBL | Immuno-Biological Laboratories<br>Co., Ltd. | Cat#27751 |
| **Software** |  |  |
| BD FACSDiva™ | www.bdbiosciences.com | Version 8.0.1 |
| FlowJo | www.flowjo.com | Version 10.7.1 |
| Pluggin: FlowAI | https://www.flowjo.com/exchange/#/ | Version 2.1 |
| Pluggin: DownSample | https://www.flowjo.com/exchange/#/ | Version 3.3 |
| GraphPad Prism | www.graphpad.com | Version 7.00 |
| Imaris | www.imaris.oxinst.com | Version 9.5.0 |
| Huygens Essential | www.svi.nl/Huygens-Software | Version 19.10 |
| Microsoft Excel | www.microsoft.com | Version 16.0 |
| Adobe Illustrator | www.adobe.com | Version CS6 (64 bit) |

